# Single-nucleus multiomics of murine gonadal cells reveals transcriptional regulatory network underlying supporting lineage differentiation

**DOI:** 10.1101/2025.02.24.639877

**Authors:** Yu-Ying Chen, Karina Rodriguez, Adriana K. Alexander, Xin Xu, Brian Papas, Martin A. Estermann, Humphrey Hung-Chang Yao

## Abstract

Sex determination of mammalian gonads hinges upon sex-specific differentiation of gonadal supporting cells: Sertoli cells in the testis and granulosa cells in the ovary. To gain insights into how supporting cells acquire their identities, we performed joint single-nucleus transcriptomics and chromatin accessibility assays on murine gonadal cells across sex determination. By contrasting sex-specific gene expression and corresponding chromatin accessibility among progenitor and differentiated cells, we discovered that sex-specific chromatin regions in supporting cells are established shortly after sex determination, accompanied by the acquisition of active histone marks. The presence of potential transcription factor binding motifs in the open chromatin regions revealed regulatory networks underlying ovary-enriched factors LEF1 and MSX1, which promote granulosa fate by inducing granulosa-specific genes such as *Foxl2* and *Fst*. Our results not only identify the gene regulatory framework underlying supporting cell sex differentiation, but also provide invaluable resources for the field.

**Teaser:** Sex-specific changes in chromatin and gene expression underly the divergent developmental fates of the bipotential gonadal primordium.

## Introduction

The gonadal primordium, also known as the genital ridge, is among the mammalian tissues that have two divergent developmental fates: testis or ovary. In mouse embryos, the bipotential gonadal primordium arises from the epithelium along the coelomic surface of the mesonephros around embryonic day (E) 10 (1, 2). As the coelomic epithelium proliferates and constitutes the somatic compartment of the gonadal primordium, it gives rise to both the supporting and interstitial cell lineages (2, 3). The supporting cell precursors, or pre-supporting cells, show minimal sex differences and remain bipotential prior to sex determination. In the XY mouse gonad, pre-supporting cells upregulate the Y-linked gene *Sry* (4, 5), which in turn induces the expression of its downstream effector *Sox9* (6). *Sox9*, along with other factors including *Fgf9* (7) and *Pgd2* (8), promotes the differentiation of pre-supporting cells into Sertoli cells (9). Conversely, in the XX mouse gonad where *Sry* is absent, pre-supporting cells give rise to pre-granulosa cells that eventually encapsulate female germ cells and form ovarian follicles perinatally (10). Following sex determination, the gonadal epithelium continues to differentiate into pre-granulosa cells, which occurs only in females but not males, contributing to the second wave of follicle formation (11, 12). The differentiation of pre-granulosa cells appears to be multi-factorial. Factors promoting pre-granulosa cells differentiation include *Wnt4* (13), *Foxl2* (14), *Fst* (15), *Runx1* (16), and the -KTS variant of *Wt1* (17). Dysregulation of these factors results in not only complete or partial sex reversal in mice but also differences of sexual development (DSD) in humans (13, 16-21). Despite the identification of these ovary-determining factors, the precise sequence of events driving pre-granulosa cell differentiation remains elusive.

Changes in chromatin accessibility are associated with supporting cell differentiation in the gonad (23, 24). In somatic progenitor cells, the majority of open chromatin regions are shared between sexes. Additionally, genes regulating sex differentiation are bivalently marked with both active trimethylation of histone H3 lysine 4 (H3K4me3) and repressive H3K27me3 histone marks at their promoters (25). By E13.5, when Sertoli and pre-granulosa fates are established, over 50% of open chromatin regions become specific to either cells. This acquisition of sex-specific accessible chromatin regions is accompanied by the loss of repressive H3K27me3 histone marks of the opposite sex (25). These observations suggest that the increase in sex-specific open chromatin regions may promote transcriptional programs unique to Sertoli or pre-granulosa cell fate.

To investigate supporting cell differentiation, we applied joint single-nucleus transcriptomics and chromatin accessibility assays (10x Genomics Multiome) in this study. This approach simultaneously profiles nuclear mRNA and DNA from individual nuclei, enabling direct linkage of gene expression and accessible chromatin profiles from the same cells. By profiling timepoints across early gonadal development, we examine how the acquisition of unique chromatin profiles relates to supporting cell fate commitment. Additionally, by mapping enriched transcription factor motifs within pre-granulosa open chromatin regions, we aim to uncover the transcriptional regulatory networks driving pre-granulosa cell differentiation.

## Results

### Single-nucleus RNA-seq of mouse embryonic gonads recapitulates cell types identified through single-cell RNA-seq

To delineate sex-specific transcriptional networks underlying somatic cell differentiation in the gonad, we performed joint single-nucleus (sn) RNA and ATAC sequencing (seq) (10x Genomics Multiome) on mouse XX and XY gonadal cells at E11.5 (initiation of sex determination), E12.5 (onset of morphological differentiation), and E13.5 (morphological dimorphism) (**Figure 1A**). The assay allowed us to obtain information on gene expression and chromatin status from the same nuclei simultaneously. After quality control and filtering (26), we profiled a total of 51,990 nuclei (XX 12,335, 13,625, 5,704, and XY 5,134, 10,246, 4,946 nuclei at E11.5, E12.5, and E13.5, respectively) (**Figure S1**). We first projected the snRNA-seq data of individual nucleus onto lower dimensional space using the Uniform Manifold Approximation and Projection (UMAP). We observed that the transcriptomes of XX and XY nuclei overlapped extensively at E11.5 and diverged apart by sex at E12.5 and E13.5 (**Figure 1B, i**). Upon clustering based on RNA data and analyzing the expression of genes known for various cell types (27), we assigned 14 major cell populations, including germ cells, epithelial cells, mesenchymal cells, pre-granulosa cells, Sertoli cells, pre-supporting cells, and supporting-like cells which constitute the future rete testis and rete ovarii (27) (**Figure 1C, i, Figure S2A**). Specifically, gonadal epithelial cells were enriched with *Upk3b* and *Aldh1a2* expression (28, 29); pre-supporting cells expressed *Wnt4* and *Runx1* and were present in both sexes at E11.5 (16); supporting-like cells were enriched with *Pax8* expression (27); while pre-granulosa and Sertoli cells were sex-specific clusters and expressed *Fst* and *Sox9,* respectively (15, 30) (**Figure 1D**). Consistent with other single cell RNA-seq datasets (27, 31), we observed a drastic decrease of the percentage of pre-supporting cells from E11.5 to E12.5 in both XX and XY gonads, accompanied by the emergence of pre-granulosa cells in E12.5 XX gonads and interstitial progenitor cells in E12.5 XY gonads (**Figure 1E, Figure S2A-E**). While we took advantage of the *Nr5a1-Cre; Rosa-tdTomato* embryos to assist with the separation of the gonad from the mesonephros, we observed a small cluster of cells from the mesonephros, i.e. *Pax2* and *Pax8* enriched mesonephric tubules (0.6%) (27). Overall, our single-nucleus RNA-seq dataset, which aligned with published single-cell RNA-seq datasets, produced reliable transcriptomic basis for further downstream chromatin analysis.

**Figure 1:**
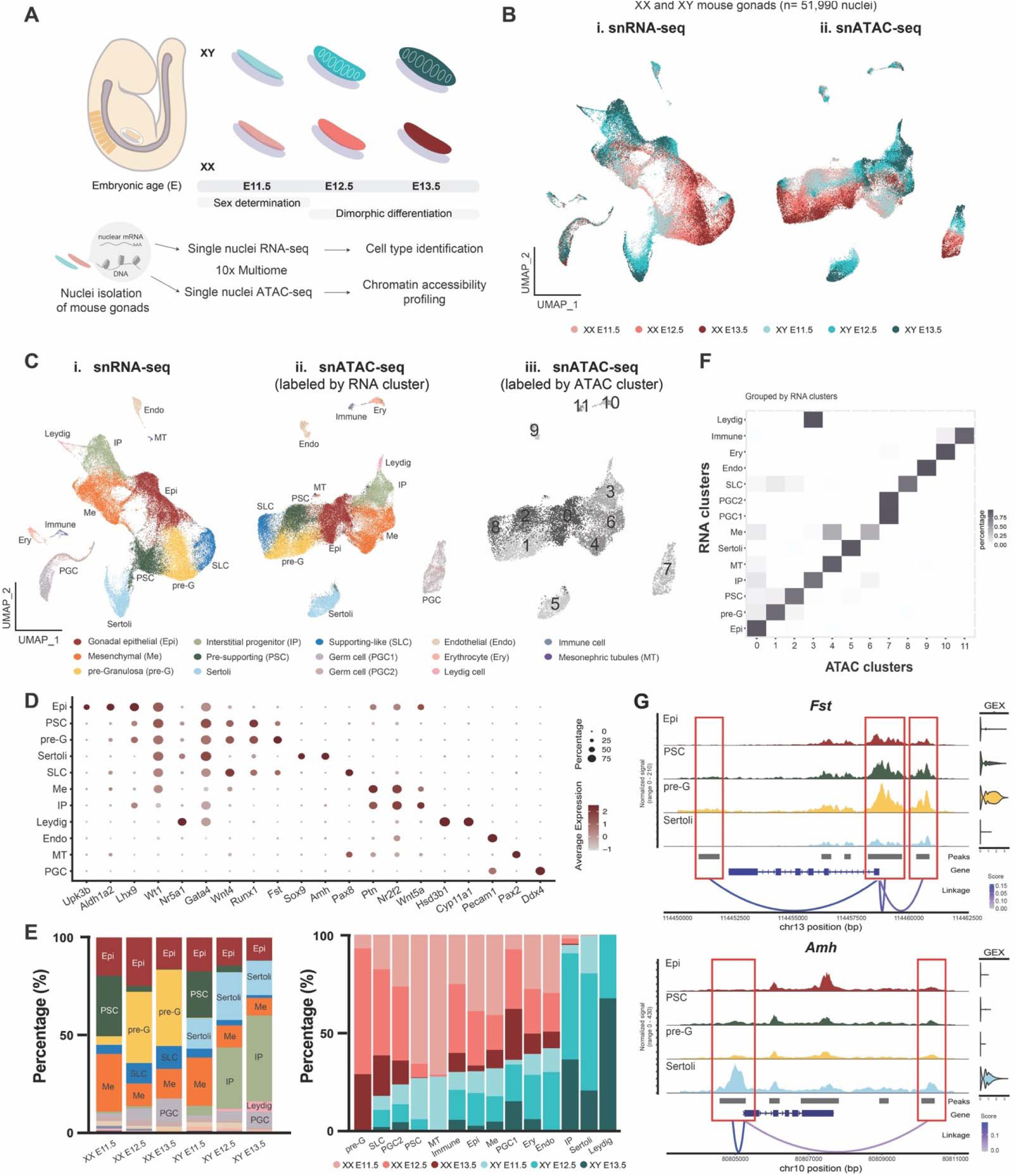
Joint single-nucleus multiomics of mouse gonadal cells during sex determination reveals highly correlated chromatin accessibility and gene expression profiles among different cell types. (A) Schematic of joint single-nucleus (sn) RNA and ATAC sequencing (seq) (10x Genomics Multiome) on mouse XX and XY gonadal cells at E11.5 (initiation of sex determination), E12.5 (onset of morphological differentiation), and E13.5 (morphological dimorphism). After nucleus isolation, nuclear mRNA was used for snRNA-seq and cell type annotation, while DNA from the same nucleus was used for snATAC-seq and chromatin accessibility profiling. (B) UMAP of snRNA-seq (i) and snATAC-seq (ii) data collected from E11.5-E13.5 XX and XY gonadal cells color coded by embryonic sex and stage. (C) UMAP of snRNA-seq (i) and snATAC-seq (ii-iii) data collected from E11.5-E13.5 XX and XY gonadal cells color coded by unbiased clustering based on either RNA (i-ii) or ATAC data (iii). Annotations for the snRNA-seq clusters are: Gonadal epithelial (Epi), Mesenchymal (Me), Pre-granulosa (pre-G), Interstitial progenitor (IP), Pre-supporting (PSC), Supporting-like cell (SLC), Germ cell (PGC), Endothelial cell (Endo), Erythrocyte (Ery), Mesonephric tubule (MT). (D) Marker gene expression for cell type annotation based on RNA clustering. (E) Percentage of cell type composition within each embryonic sex and stage (left), and percentage of sex and stage composition within each cell type (right). (F) Heatmap of cluster distribution of cells based on ATAC clustering (x-axis) and RNA-clustering (y-axis), as grouped by individual RNA cluster (the sum of each row is 1). (G) Peak-gene linkage plots of *Fst* (top) and *Amh* (bottom) in epithelial (EPI), pre-supporting (PSC), pre-granulosa (pre-G), and Sertoli cells of combined sex and developmental stages. The red boxes denote called peaks (grey bars) that are linked to gene expression (GEX). Linkage score represents the level of association between chromatin accessibility and gene expression.

### Chromatin accessibility and gene expression profiles of gonadal somatic cells are highly correlated

Following cell type annotation via snRNA-seq, we performed peak calling using MACS2 (32) on individual RNA clusters to capture cell type-specific ATAC peaks. Similar to snRNA-seq data, UMAP of snATAC-seq data showed overlapping of XX and XY nuclei at E11.5 and divergence by sex at E12.5 and E13.5 (**Figure 1B, ii**). When color-coded by cell types, Sertoli cells exhibited complete separation from other somatic cells, reflecting global chromatin remodeling that differentiate Sertoli from precursor cells as early as E11.5 (**Figure 1C, ii**). Upon clustering based on ATAC data, we observed that chromatin states generally correlated with gene expression among major cell types (**Figure 1C, ii, iii, & 1F**). Despite the correlation, small proportions of pre-supporting cells and supporting-like cells shared the same ATAC cluster with pre-granulosa cells, indicating a closer similarity in their chromatin landscape (**Figure 1F**). In addition, mesenchymal cells encompassed various chromatin states, while Leydig cells shared similar chromatin profile with interstitial progenitors (**Figure 1F**). Finally, to correlate accessible chromatin peaks to gene expression, we performed linkage analysis (33) on the whole dataset to obtain peaks and genes that were statistically significantly associated with one another within transcription start site (TSS) ± 500Kb (linkage distance) as compared across cell types (**Data S1**). Using canonical pre-granulosa cell marker *Fst* and Sertoli marker *Amh* as examples, we identified that the promoter peak and the peaks right upstream and downstream of *Fst* were positively correlated to *Fst* expression (**Figure 1G**). These peaks were accessible in epithelial, pre-supporting, and pre-granulosa cells, but not in Sertoli cells. Whereas the promoter peak and a peak downstream of *Amh* were positively correlated to *Amh* expression (**Figure 1G**). These two peaks were only accessible in Sertoli cells but not in pre-granulosa and precursor cell types. In contrast, the intragenic peaks of *Amh* were not associated to *Amh* expression as they were also present in other cell types (**Figure 1G**). Together, our analyses revealed the dynamic changes in chromatin modulation during gonadal cell differentiation, and provided an opportunity to investigate how changes in chromatin accessibility across cell types and developmental timepoints are associated with changes in gene expression.

### Sex differences in chromatin accessibility between Sertoli and pre-granulosa cells augment during E11.5 to E12.5 transition

To understand how chromatin profiles of gonadal somatic cells change during sex determination, we compared accessible chromatin regions (differential accessible peaks, DAPs) between sexes across cell types and developmental timepoints (**Figure 2A, Data S2**). The greatest number of DAPs and the most changes in DAP numbers were observed between Sertoli and pre-granulosa cells from E11.5 to E12.5, followed by epithelial cells (**Figure 2A**). The XY mesenchymal cells also showed drastic changes in sex-specific chromatin pattern from E11.5 to E12.5. On the other hand, pre-supporting cells, supporting-like cells, and endothelial cells showed minimal sex differences in terms of accessible chromatin regions (**Figure 2A**). Sertoli cells displayed nearly three times (E11.5) and twice (E12.5) as many DAPs than pre-granulosa cells, reflecting the delayed timeline in pre-granulosa differentiation (11). Upon annotating the genomic region of all DAPs, we found that the majority of the Sertoli and pre-granulosa specific DAPs at E11.5 were located within distal intergenic regions and other intronic regions, followed by the promoter regions (**Figure 2B**).

**Figure 2:**
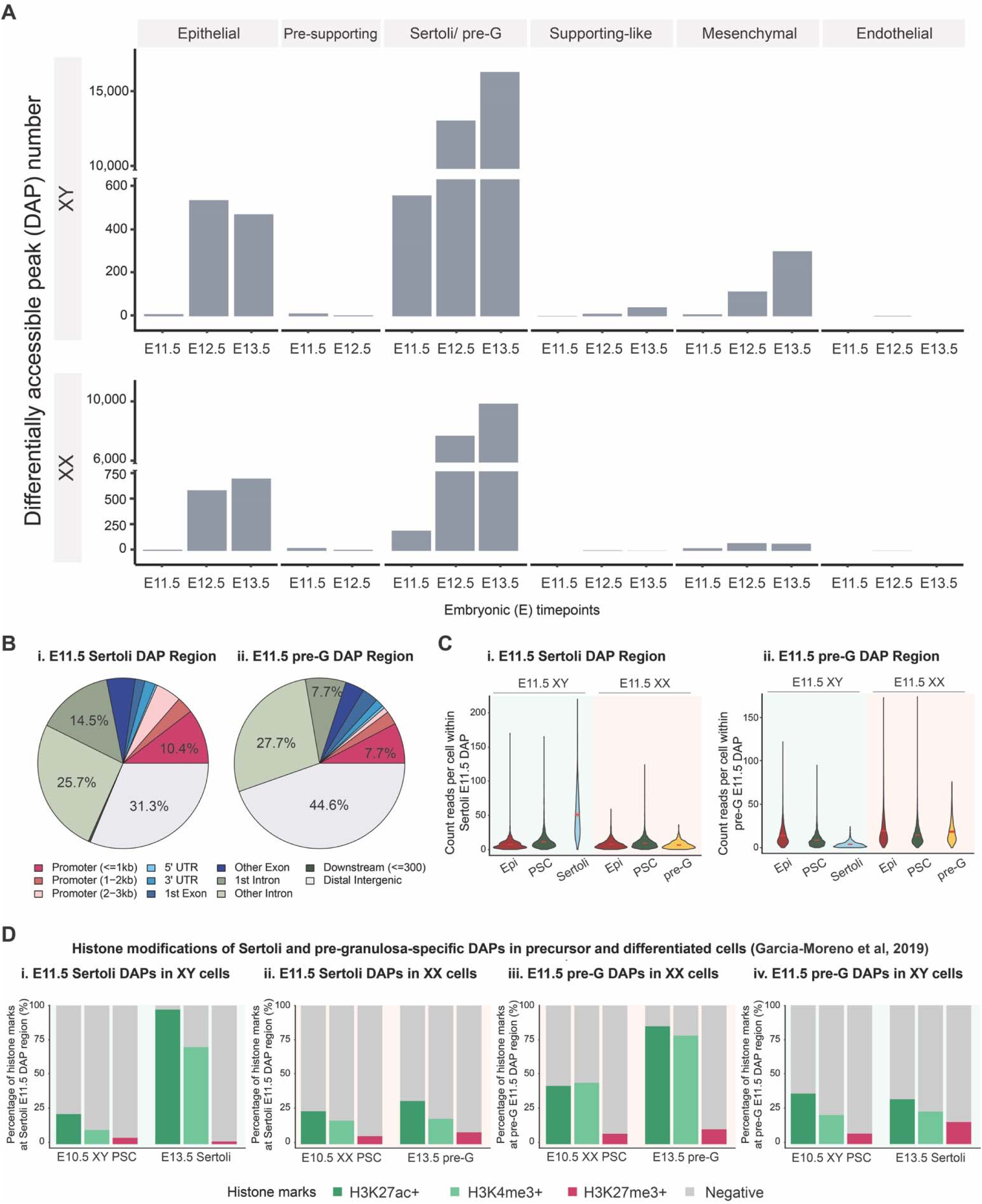
Sertoli and pre-granulosa cells undergo active chromatin and epigenetic changes during sex differentiation. (A) Numbers of differentially accessible peaks (DAPs) between sexes across cell types and developmental timepoints. (B) Peak annotation within E11.5 Sertoli DAPs (i) and E11.5 pre-granulosa cell DAPs (ii) (C) Chromatin accessibility, measured by average count reads per cell within E11.5 Sertoli DAPs (i) and pre-granulosa E11.5 DAPs (ii) of E11.5 XX and XY epithelial, pre-supporting cells, Sertoli, and pre-granulosa cells. (D) Percentage of H3K27ac, H3K4me3, and H3K27me3-positive chromatin regions (obtained from published ChIP-seq datasets, Garcia-Moreno et al, 2019) that overlap with E11.5 Sertoli DAPs (i, ii) and E11.5 pre-granulosa DAPs (iii, iv) in XY (I, iv) and XX (ii, iii) pre-supporting, Sertoli or pre-granulosa cells.

To explore how Sertoli and pre-granulosa cells acquired sex-specific DAPs by E11.5, we measured chromatin accessibility within E11.5 Sertoli DAPs (537 unique peaks) and pre-granulosa E11.5 DAPs (195 unique peaks) in the precursor cell types, namely E11.5 XX and XY epithelial and pre-supporting cells (**Figure 2C**). We found that the average accessibility within E11.5 Sertoli DAPs was significantly higher in Sertoli than in pre-granulosa cells as expected. In XY gonads, the average accessibility increased slightly in pre-supporting cells, then further increased drastically in Sertoli cells as compared to XY pre-supporting and epithelial cells. This increase was not observed in pre-granulosa cells (**Figure 2C, i**). On the other hand, the average accessibility within E11.5 pre-granulosa DAPs was significantly higher in pre-granulosa than in Sertoli cells, albeit at a lower degree than the difference within E11.5 Sertoli DAPs (**Figure 2C, ii**). In XX gonads, the average accessibility decreased slightly in pre-supporting cells, then increased slightly in pre-granulosa cells as compared to XX pre-supporting cells. In XY gonads, average accessibility decreased slightly in pre-supporting, then decreased drastically in Sertoli cells as compared to XY epithelial cells (**Figure 2C, ii**). When examining Sertoli and pre-granulosa E12.5 DAPs (12,991 and 7,727 unique peaks respectively), we observed a similar trend within Sertoli DAPs, while partial pre-granulosa cells exhibited highest accessibility level within pre-granulosa DAPs (**Figure S3A, B**). We further examined changes in the accessibility of E13.5 pre-granulosa-specific chromatin regions in pre-granulosa cells. We observed a continuous increase in accessibility throughout timepoints in pre-granulosa cells. This suggested that the acquisition of pre-granulosa DAPs occurred not only passively, through Sertoli cells decreasing accessibility in these regions, but also actively, with pre-granulosa cells gaining chromatin accessibility (**Figure S3C**). These results indicated that Sertoli and pre-granulosa cells undergo active chromatin remodeling during sex differentiation, with Sertoli cells exhibiting a greater change. During Sertoli cell differentiation, Sertoli DAPs become more accessible while regions overlapping pre-granulosa DAPs become less accessible. Whereas during pre-granulosa differentiation, chromatin regions overlapping Sertoli and pre-granulosa DAPs alter in much less extend as compared to Sertoli cells.

### Sertoli and pre-granulosa specific chromatin regions acquire distinctive active and repressive histone marks during sex differentiation

Sertoli and pre-granulosa-enriched genes are bivalent at their promoter regions before sex determination, marked by both active H3K4me3 and repressive H3K27me3 histone marks (25). During sex determination, genes promoting one fate lose the repressive H3K27me3 marks, while genes promoting the alternate fate remain bivalent (25). To explore how the acquisition of Sertoli and pre-granulosa-specific DAPs correlates with changes in histone marks, we examined the status of histone modifications within these DAPs in precursor and differentiated cells. We analyzed published ChIP-seq data for H3K4me3, H3K27me3, and the active acetylation of histone H3 lysine 27 (H3K27ac) from E10.5 purified mouse bipotential progenitor cells (referred to as pre-supporting cells or PSCs hereafter) and E13.5 Sertoli and pre-granulosa cells (23, 25) (**Figure 2D**). We observed a drastic increase in active H3K4me3 and H3K27ac histone marks within Sertoli-specific DAPs (537 unique peaks) during Sertoli cell differentiation. The proportion of H3K4me3-positive Sertoli-specific DAPs rose from 10% in E10.5 pre-supporting to 70% in E13.5 Sertoli cells, while H3K27ac-positive Sertoli-specific DAPs increased from 22% in E10.5 pre-supporting to over 97% in E13.5 Sertoli cells (**Figure 2D, i**). In contrast, the levels of H3K4me3 and H3K27ac-positive Sertoli-specific DAPs showed minimal changes during pre-granulosa differentiation (**Figure 2D, ii**). For pre-granulosa-specific DAPs (195 unique peaks), E10.5 XX pre-supporting cells displayed higher levels of H3K4me3 and H3K27ac marks (45% and 42%, respectively) (**Figure 2D, iii**) as compared to Sertoli-specific DAPs in E10.5 XY pre-supporting cells (**Figure 2D, i**). Following pre-granulosa cell differentiation, the proportion of H3K4me3 and H3K27ac-positive pre-granulosa-specific DAPs increased to 78% and 85%, respectively (**Figure 2D, iii**). H3K4me3 and H3K27ac-positive pre-granulosa-specific DAPs remained at a similar level during Sertoli cell differentiation (**Figure 2D, iv**).

For the repressive H3K27me3 modifications, we did not observe significant changes overall. Within Sertoli-specific DAPs, H3K27me3 level decreased from 5% in E10.5 XY pre-supporting to 2% in E13.5 Sertoli cells (**Figure 2D, i**), whereas H3K27me3 level increased slightly from 6% in E10.5 XX pre-supporting to 9% in E13.5 pre-granulosa (**Figure 2D, ii**). Within pre-granulosa-specific DAPs, H3K27me3 level increased both in Sertoli (8% to 16%) (**Figure 2D, iv**) and pre-granulosa cells (8% to 11%) (**Figure 2D, iii**). Upon overlapping H3K4me3 and H3K27me3 peaks, we found that only 8 out of 537 E11.5 Sertoli cell-specific DAPs were bivalent for both H3K4me3 and H3K27me3 marks in E10.5 XY pre-supporting cells. Among them, three were linked to genes *Myh14*, *Rpl13a*, and *Gramd2*. In the case of pre-granulosa cells, 13 out of 195 E11.5 DAPs were bivalent for both histone marks in E10.5 XX pre-supporting cells. Among them, 4 were linked to genes *Ereg*, *Cbln1*, *Sp5*, and *Rpl12* (**Table S1**). Similar histone methylation and acetylation patterns were observed examining Sertoli and pre-granulosa E13.5 DAPs (16,235 and 9,886 unique peaks, respectively) in pre-supporting and differentiated cells (**Figure S4**). Collectively, these results suggested that the acquisition of sex-specific accessible chromatin regions is accompanied by the deposition of active H3K4me3 and H3K27ac histone marks. Sertoli cells exhibited a 4- to 7-fold increase, while pre-granulosa cells showed a 2-fold increase, reflecting broader changes in chromatin accessibility in Sertoli cells compared to pre-granulosa cells (**Figure 1C, ii**).

### Increase in Sertoli and pre-granulosa-specific gene expression from E11.5 to E12.5 is associated with changes in chromatin accessibility

As we observed active changes in chromatin landscape during supporting cell differentiation, we next explored the association of chromatin accessibility with gene expression. We first obtained differentially expressed genes (DEGs) between sexes across cell types and developmental timepoints. We then categorized DEGs that were 1) positively or negatively linked to any DAPs, 2) positively or negatively linked to chromatin accessible peaks that were not DAPs between sexes, or 3) not linked to any peaks at all (**Figure 3A, Data S3**). For the first category, we defined DEGs with at least one linked DAP as “DEG with linked DAP”. An example of this category was *Amh,* a Sertoli DEG. Expression of *Amh* was linked to the *Amh* promoter peak, which was differentially accessible between Sertoli and pre-granulosa cells (**Figure 3A, i**). In the second category, when a DEG was linked to a chromatin accessible peak, however the peak itself was not a DAP, we classified it “DEG with linked non-DAP" (**Figure 3A, ii**). For instance, the scramblase-encoding gene *Xkr4* was identified as a Sertoli DEG compared to pre-granulosa cells. The expression of *Xkr4* was linked to its promoter peak, as analyzing across the entire dataset, this promoter peak was accessible and the gene was expressed in Sertoli cells, whereas in another cell type, such as immune cells, the promoter region of *Xkr4* was not accessible and the gene was not expressed (**Figure 3A, ii and Figure S5A**). Although *Xkr4* was a Sertoli DEG, its promoter peak was not a DAP between Sertoli and pre-granulosa cells, rendering it a “DEG with linked non-DAP" between the two cell types (**Figure 3A, ii**). In the third category, an example was the Potassium voltage-gated channel *Kcnq5*, a Sertoli DEG. Its expression was not associated with any present peaks within the TSS ± 500kb linkage distance across cell types (**Figure S5B**), therefore it was assigned as “DEG with no linked-peak” (**Figure 3A, iii**).

**Figure 3:**
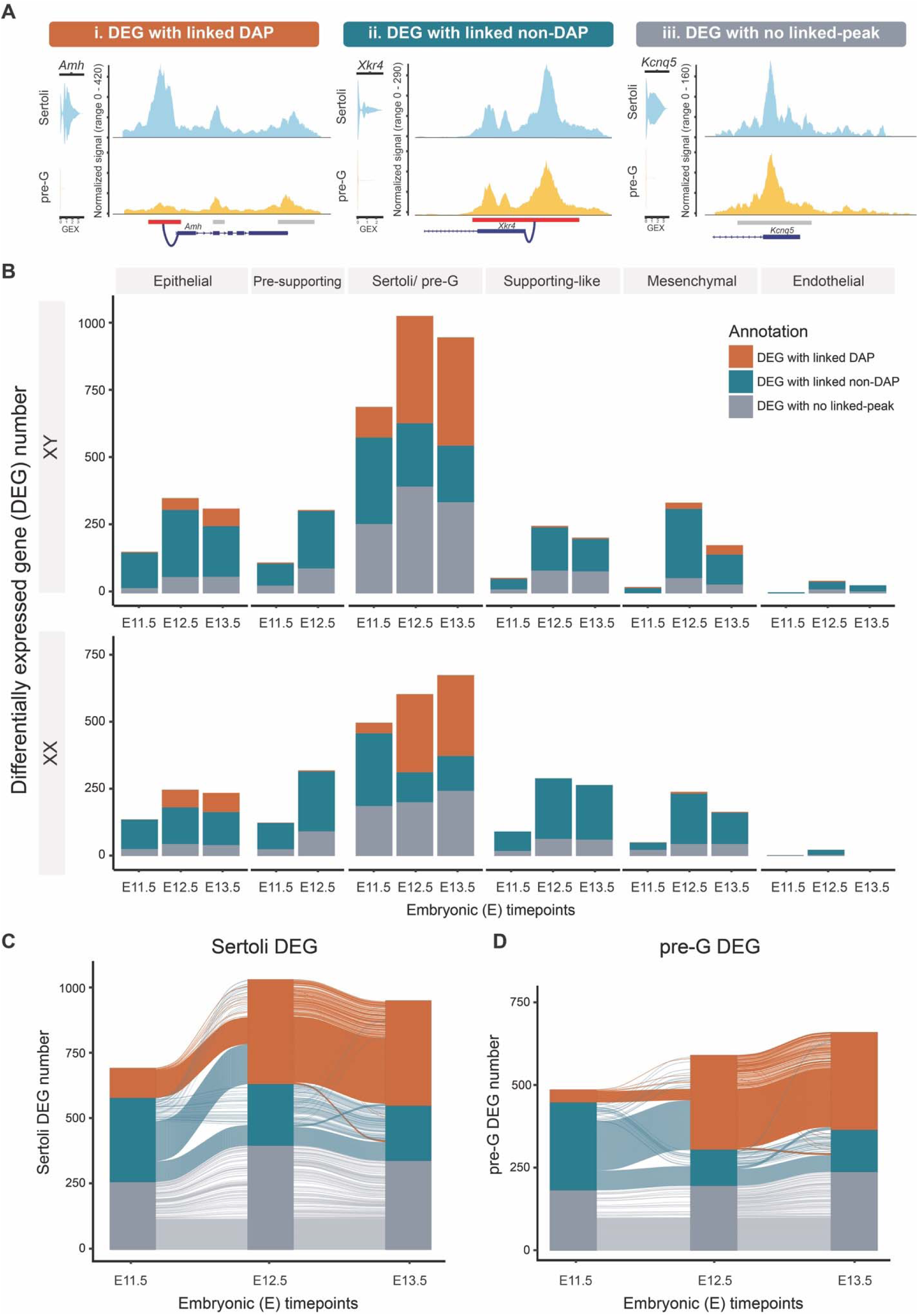
The association between Sertoli and pre-granulosa-specific gene expression and differential chromatin pattern increases during sex determination. (A) Categorization and examples of DEGs with linked differentially accessible peaks (DAPs) between sexes (i), linked to chromatin accessible peaks that were not DAPs between sexes (ii), or not linked to any peaks (iii). (B) Numbers of DEGs between sexes across cell types and developmental timepoints, annotated with linkage categorization. (C, D) Changes in the annotation of three DEG categories in Sertoli (C) and pre-granulosa cells (D) over developmental timepoints visualized with Sankey diagram.

Based on this analysis, we observed that Sertoli and pre-granulosa cells exhibited the highest number of DEGs across all time points from E11.5 to E13.5 (**Figure 3B**). Among these, “DEG with no linked-peak” accounted for more than 30% of DEGs, suggesting the possibility that gene expression in these cases is regulated by sex-specific transcription factor binding in accessible chromatin regions, or through potential distal enhancer action beyond the linkage distance. Strikingly, the proportion of “DEG with linked DAP” increased from 16% to 38% in Sertoli and 8% to 48% in pre-granulosa cells between E11.5 and E12.5 (**Figure 3B**). In epithelial, pre-supporting, supporting-like, and mesenchymal cells, an increase in DEG number was observed between E11.5 and E12.5, with the majority of DEGs linked to non-DAPs (**Figure 3B**).

To explore the dynamics of peak-gene association across timepoints, we visualized changes in the annotation of three DEG categories in Sertoli and pre-granulosa cells (**Figure 3C, D**). We observed that “DEG with linked DAP” at E11.5 retained the same annotation across timepoints. In contrast, a large proportion of “DEGs with linked non-DAPs" at E11.5 converted to “DEG with linked DAP” at E12.5, accounting for 53% in Sertoli cells and 60% in pre-granulosa cells (teal to orange transition in **Figure 3C, D**). Although the number of DEGs with no linked-peaks increased slightly, it was not due to conversion of other two categories. The distribution of the three DEG categories stabilized from E12.5 to E13.5 (**Figure 3C, D**). These results indicated that Sertoli and pre-granulosa cells undergo active chromatin remodeling, particularly during the transitioning from E11.5 to E12.5. The observed increase in sex-specific gene expression that were linked to changes in chromatin status suggested potential modulation through transcription factor (TF) binding during early differentiation.

### Motif analysis reveals transcription factor network underlying Sertoli differentiation

Given the dynamic changes in sex-specific chromatin accessible regions (DAPs) associated with DEGs during supporting cell differentiation from E11.5 to E12.5, we hypothesized that transcription factor binding motifs enriched within E11.5 DAPs could be responsible for early dimorphic differentiation, while those enriched within E12.5 DAPs represent the downstream, secondary factors. These transcription factors together constitute the transcriptional regulatory network underlying supporting cell differentiation. To explore such network driving Sertoli cell differentiation, we performed transcription factor binding motif analysis using the workflow outlined in **Figure 4A**. First, we performed transcription factor motif enrichment analysis on Sertoli DAPs derived from “DEG with linked DAP”. Second, we overlapped the enriched motifs and Sertoli cell DEGs, and identified candidate motifs with enriched transcription factors as DEGs. Finally, we performed motif scan analysis (34) to determine transcription factor motif positions within Sertoli DAPs, and established the association of putative transcription factors with their downstream target genes (**Figure 4A**). Transcription factor binding motif analysis on E11.5 Sertoli “DEG with linked-DAP" (**Figure 3C**, E11.5 orange) revealed SOX transcription factors, including SOX10, SOX13, SOX4, SOX6, and SOX9 as early transcription factors regulating Sertoli differentiation (**Figure 4B**). Gene expression of these SOX factors showed partial expression in E11.5 XX and XY epithelial cells, low level of expression in XY pre-supporting cells, and an upregulation in Sertoli cells. The expression was absent in XX pre-supporting and pre-granulosa cells (**Figure 4C**). Upon locating SOX binding motifs within Sertoli DAPs and associating these DAPs with expression of other genes, we found that SOX motifs-enriched DAPs were positively linked to the expression of Sertoli enriched genes, including *Sox9*, *Dmrt1*, *Inhbb*, *Fstl4*, and *Cyp26b1*. Conversely, SOX motifs-enriched DAPs were negatively linked to genes absent in Sertoli cells, such as *Gli2*, *Tcf4*, and *Rbms3* (**Figure 4D**). Among the three negatively associated genes, *Gli2* and *Tcf4* showed higher expression in pre-granulosa cells (**Figure 4D**).

**Figure 4:**
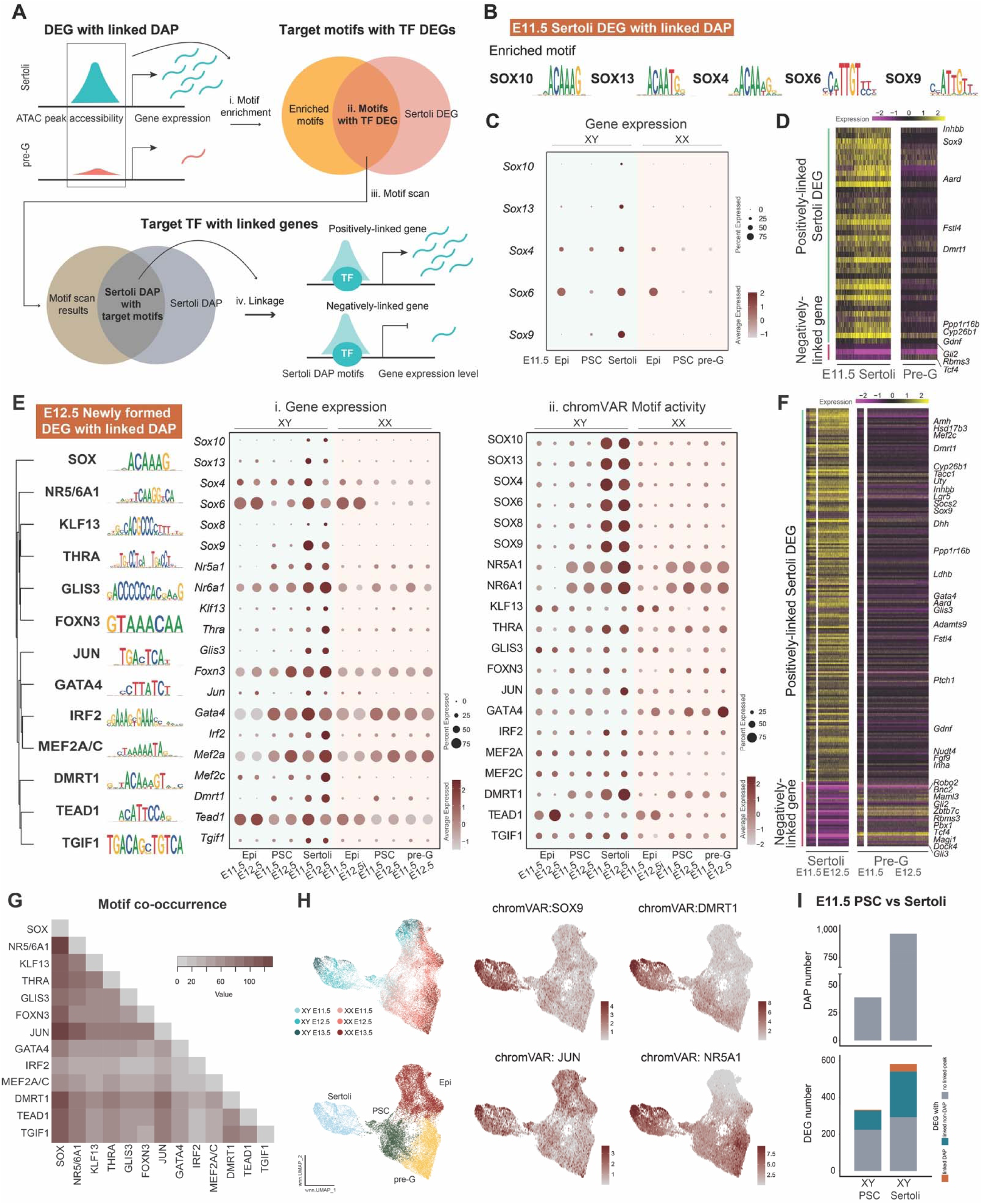
The transcription regulatory network underlying Sertoli cell differentiation. (A) Workflow of transcription factor binding motif analysis: Motif enrichment was first performed on Sertoli DAPs derived from “DEG with linked DAP” (i). Candidate motifs with expressed transcription factors were then identified by overlapping the motifs with Sertoli DEGs (ii). Following, motif scan was performed to determine motif position within Sertoli DAPs (iii). Lastly, downstream target genes of the motifs were identified based on the linkage analysis (iv). (B) Identified motifs enriched within “E11.5 Sertoli DEGs with linked DAP”, with position weight matrix of the corresponding motifs. (C) Gene expression of the identified TFs in E11.5 XX and XY epithelial (Epi), pre-supporting (PSC), Sertoli, and pre-granulosa cells. (D) Gene expression heatmap of the positively (green) and negatively (magenta)-linked downstream target genes of the identified TFs in E11.5 Sertoli and pre-granulosa cells. (E) Identified motifs enriched within “E12.5 Sertoli DEGs with linked DAP”, with position weight matrix of the corresponding motifs. Motifs were clustered based on downstream target gene enrichment. Gene expression (i) and chromVAR motif activity (ii) of the identified TFs in E11.5 and E12.5 XX and XY epithelial, pre-supporting, Sertoli, and pre-granulosa cells were shown. (F) Gene expression heatmap of the positively (green) and negatively (magenta)-linked downstream target genes of the identified TFs in E11.5 and E12.5 Sertoli and pre-granulosa cells. (G) Heatmap of motif co-occurrence based on the incidence (value) of motif sites enriched among “E12.5 Sertoli DEGs with linked DAP” associating with the same downstream target genes. (H) Feature plots of chromVAR motif activity of identified motifs on joint RNA and ATAC UMAP composed of epithelial, pre-supporting, Sertoli, and pre-granulosa cells from both sexes throughout developmental timepoints. (I) Numbers of differentially accessible peaks (DAPs) (top) and annotated DEGs (bottom) between E11.5 pre-supporting cells and Sertoli cells.

We next identified the downstream, secondary transcription factors by searching the enriched motifs within Sertoli-specific chromatin regions that were associated with Sertoli DEGs at E12.5 (**Figure 3C**, E12.5 orange). Specifically, we analyzed two groups of accessible regions: those that converted from “DEGs with linked non-DAP" to “DEGs with linked DAP” (teal to orange transition from E11.5 to E12.5 in **Figure 3C**), as well as “DEG with linked DAP” that appeared *de novo* at E12.5 (excluding orange to orange transition from E11.5 to E12.5 in **Figure 3C**). We named these two groups “E12.5 newly formed DEG with linked DAP”. Transcription factor binding motif analysis on DAPs of these two groups revealed many more motifs with enriched and expressed TFs such as DMRT1, NR5A1, GATA4, NR6A1, JUN, and IRF2 in addition to SOX factors (**Figure 4E**). Different gene expression patterns of these transcription factors were observed: factors including *Sox4*, *Sox6*, *Nr6a1*, and *Tead1* showed expression starting in XY epithelial cells and increased expression in pre-supporting cells and Sertoli cells. On the other hand, *Sox9*, *Nr5a1*, *Foxn3*, *Gata4*, and *Mef2a* showed expression starting in XY pre-supporting cells and continued in Sertoli cells. Finally, *Sox10*, *Sox13*, *Sox8*, *Klf13*, *Thra*, *Glis3*, *Irf2*, and *Tgif1* showed increased expression only in Sertoli cells (**Figure 4E, i**). Gene expression of most transcription factors was absent in XX cells with small exceptions: *Sox6* and *Tead1* were observed in XX epithelial cells but not in pre-supporting or pre-granulosa cells; *Gata4* and *Mef2a* were observed in pre-supporting and pre-granulosa cells, despite in lower level than Sertoli cells (**Figure 4E, i**). These transcription factors were positively associated with the expression of Sertoli DEGs such as *Amh*, *Lgr5*, *Dhh*, *Ptch1*, *Inha*, while negatively associated with the expression of *Bnc2*, *Zbtb7c*, *Pbx1*, *Tcf4*, and *Gli3* (**Figure 4F**). Many of the negatively associated genes were expressed in higher level in pre-granulosa cells (**Figure 4F**). To investigate whether these TFs can regulate the same target genes, we generated a motif co-occurrence matrix by calculating the incidence of TFs within motif sites that were associated with the same target genes. The highest co-occurrence of TF binding sites was found in SOX with NR5/6A1 and JUN, followed by SOX with DMRT1 and NR5/6A1 with JUN (**Figure 4G**).

To provide further evidence for the identification of these putative transcription factors, we applied chromVAR (35), a program that infers transcription factor-associated chromatin accessibility, or motif activity, from single-cell ATAC-seq data. We found that SOX TFs, DMRT1, THRA, and JUN showed high motif activity in Sertoli cells (**Figure 4E, ii**). ChromVAR also predicted that NR5A1 and NR6A1 were active motifs in both XX and XY pre-supporting cells but with enhanced activity in E12.5 Sertoli cells (**Figure 4E, ii**). To visualize the motif activity at the single-cell level, we plotted chromVAR motif activity score on a joint RNA and ATAC UMAP of epithelial, pre-supporting, Sertoli, and pre-granulosa cells (**Figure 4H**). The SOX9 motif activity increased in Sertoli cells, while DMRT1 activity exhibited a gradual increase in pre-supporting cells and peaked in Sertoli cells. The motif activity for JUN showed broad distribution, with its highest level observed in Sertoli cells. Lastly, NR5A1 motif activity increased in pre-supporting cells and peaked in both Sertoli and pre-granulosa cells (**Figure 4H**). The observation that some motifs showed increased activity in pre-supporting cells before Sertoli cells, while others peaked directly in Sertoli cells prompted us to compare the two cell types directly. Comparing E11.5 XY pre-supporting cells with E11.5 Sertoli cells, we found that Sertoli cells underwent active chromatin remodeling and acquired Sertoli specific chromatin pattern during cell differentiation. Sertoli cells exhibited a 24-fold increase in DAPs and a 1.8-fold increase in DEGs compared to pre-supporting cells (**Figure 4I**). Among DEGs linked to DAPs, genes including *Msx1* and *Emx2* were downregulated in pre-supporting cells, whereas *Sox9*, *Sox10*, *Lgr5*, and *Amh* displayed increased expression in Sertoli cells (**Data S4, 5**). These genes likely were regulated from changes on the chromatin level during pre-supporting to Sertoli transition.

### Motif analysis reveals LEF1 and MSX1 as initial TFs underlying pre-granulosa differentiation

Having established a pipeline that identified known transcriptional regulators of Sertoli cell differentiation as proof of concept, we applied the same approach to pre-granulosa cells. Using this approach, we overlapped the enriched transcription factor motifs within pre-granulosa DAPs and pre-granulosa DEGs to identify putative TFs. Next, we determined the positions of TF motifs within pre-granulosa DAPs and associated them with positively or negatively regulated downstream target genes (**Figure 5A**). Such analysis on E11.5 pre-granulosa “DEGs with linked-DAPs" revealed LEF1 and MSX1 as TFs with enriched motifs (**Figure 5B**). *Lef1* expression was observed in E11.5 XX epithelial, pre-supporting, and pre-granulosa cells but decreased in E11.5 XY epithelial and pre-supporting cells, and absent in Sertoli cells. *Msx1* was expressed in both pre-supporting cells of both sexes and remained expressed in pre-granulosa but not in Sertoli cells (**Figure 5C**). Linkage analysis of gene expression and chromatin accessibility further identified pre-granulosa DEGs, including *Msx1* and *Klf7*, as potential direct targets of LEF1, while *Tcf4* expression could be regulated by MSX1 (**Figure 5B, D**). Conversely, *Tacc1*, *Socs2*, and *Nudt4* were identified as genes negatively linked with LEF and MSX1, with their expression absent in pre-granulosa but present in Sertoli cells (**Figure 5D**).

**Figure 5:**
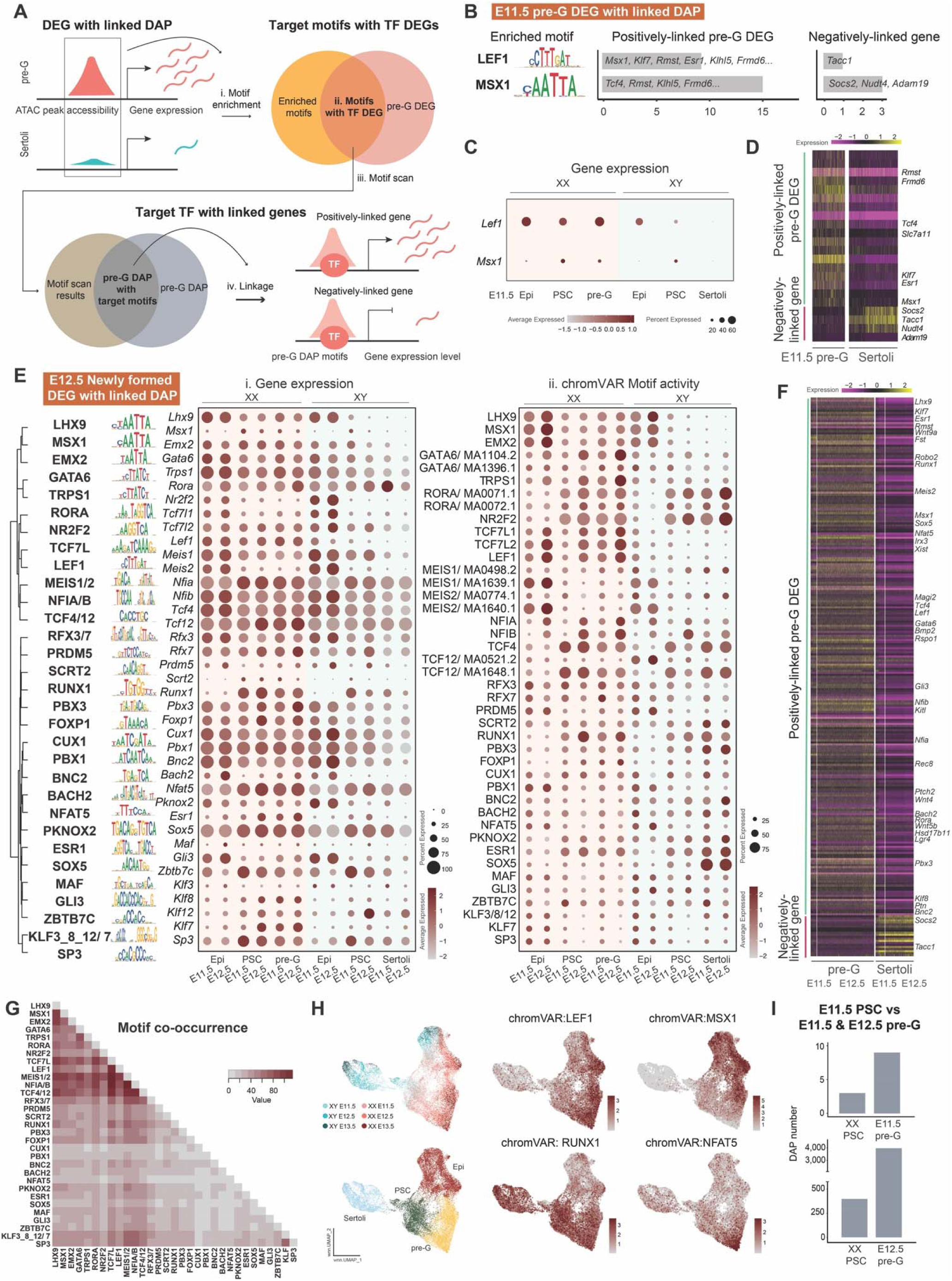
The transcription regulatory network underlying pre-granulosa differentiation. (A) Workflow of transcription factor binding motif analysis: Motif enrichment was first performed on pre-granulosa DAPs derived from “DEG with linked DAP” (i). Candidate motifs with expressed transcription factors were then identified by overlapping the motifs with pre-granulosa DEGs (ii). Following, motif scan was performed to determine motif position within pre-granulosa DAPs (iii). Lastly, downstream target genes of the motifs were identified based on the linkage analysis (iv). (B) Identified motifs enriched within “E11.5 pre-granulosa DEGs with linked DAP”, with position weight matrix of the corresponding motifs. (C) Gene expression of the identified TFs in E11.5 XX and XY epithelial (Epi), pre-supporting (PSC), pre-granulosa, and Sertoli cells. (D) Gene expression heatmap of the positively (green) and negatively (magenta)-linked downstream target genes of the identified TFs in E11.5 pre-granulosa and Sertoli cells. (E) Identified motifs enriched within “E12.5 pre-granulosa DEGs with linked DAP”, with position weight matrix of the corresponding motifs. Motifs were clustered based on downstream target gene enrichment. Gene expression (i) and chromVAR motif activity (ii) of the identified TFs in E11.5 and E12.5 XX and XY epithelial, pre-supporting, pre-granulosa, and Sertoli cells were shown. (F) Gene expression heatmap of the positively (green) and negatively (magenta)-linked downstream target genes of the identified TFs in E11.5 and E12.5 pre-granulosa and Sertoli cells. (G) Heatmap of motif co-occurrence based on the incidence (value) of motif sites enriched among “E12.5 pre-granulosa DEGs with linked DAP” associating with the same downstream target genes. (H) Feature plots of chromVAR motif activity of identified motifs on joint RNA and ATAC UMAP composed of epithelial, pre-supporting, Sertoli, and pre-granulosa cells from both sexes throughout developmental timepoints. (I) Number of differentially accessible peaks (DAPs) between E11.5 pre-supporting cells and E11.5 pre-granulosa cells (top), and between E11.5 pre-supporting cells and E12.5 pre-granulosa cells (bottom).

We next investigated enriched motifs within the newly formed DEGs with linked DAP regions in E12.5 pre-granulosa cells to identify downstream, secondary TFs regulating pre-granulosa differentiation. These regions include DEGs that converted from “with linked non-DAP” at E11.5 to become linked to at least one DAP at E12.5 (teal to orange in **Figure 3B**), as well as “DEGs with linked DAPs” that appeared *de novo* at E12.5 (orange in **Figure 3B**). Motif analysis of these regions revealed many more enriched and expressed TFs beyond LEF1 and MSX1 (**Figure 5E**). Among these motifs, BNC2, ZBTB7C, PBX1, TCF4, and GLI3 were negatively linked targets of Sertoli motifs (**Figure 4F**). Unlike most Sertoli TFs that increased gene expression in XY cells and were absent in XX cells, the majority of pre-granulosa TFs were expressed in XY precursor cells and downregulated in Sertoli cells (**Figure 5E**). We observed expression patterns that remained consistent across XX epithelial, pre-supporting, and pre-granulosa cells, including *Emx2*, *Trps1*, *Lef1*, and *Pbx1*. Factors including *Msx1*, *Scrt2*, *Runx1*, *Klf7* and *Sp3* were upregulated in pre-supporting cells. Notably, no factors showed increased expression exclusively in pre-granulosa cells (**Figure 5E**). Positively linked pre-granulosa DEGs associated with these TFs included several of the identified TFs themselves, along with *Fst*, *Irx3*, *Kitl*, *Wnt4*, and *Lgr4*. Conversely, negatively linked genes, such as *Socs2* and *Tacc1*, were expressed in Sertoli but absent in pre-granulosa cells (**Figure 5F**). To determine whether these TFs regulate the same target genes, we generated a motif co-occurrence matrix by calculating the incidence of TFs within motif sites that were associated with the same target genes. The analysis revealed a high co-occurrence rate for TCF7L with LEF1, MEIS1/2, NFIA/B, and TCF4/12, followed by LHX9 with MSX1, EMX2, TCF7L, MEIS1/2, and TCF4/12 (**Figure 5G**), suggesting these factors may regulate target gene expression synergistically.

When we computed motif activity using chromVAR, we observed that several pre-granulosa motifs showed high activity in XX precursor cell types, compared to the Sertoli motifs that exhibited increased activity specifically in Sertoli cells (**Figure 5E, 4E**). We also observed that several motifs, including RORA, NR2F2, TCF4/12, PBX3, and SOX5, were predicted to be active in Sertoli cells despite the absence of corresponding gene expression (**Figure 5E**). Visualizing the motif activity on a UMAP of epithelial, pre-supporting, Sertoli, and pre-granulosa cells revealed distinct patterns: LEF1 and MSX1 showed high activity in epithelial and pre-supporting cells, RUNX1 exhibited increased activity in pre-supporting cells, whereas NFAT5 displayed heightened motif activity in pre-granulosa cells (**Figure 5H**).

To further investigate the transition from pre-supporting cells to pre-granulosa cells, we examined the changes of DAP numbers between pre-supporting cells and pre-granulosa cells at E11.5 and E12.5. We found minimal differences in chromatin patterns between these two cell types at E11.5 compared to those at E12.5 (**Figure 5I**). DAPs acquired in E11.5 pre-granulosa cells were associated to *Klf7* and *Sulf1* (**Data S6**). In contrast, a comparison between E11.5 pre-supporting cells and E12.5 pre-granulosa cells revealed significant increased acquisition of pre-granulosa-specific chromatin regions (**Figure 5I**). These included chromatin regions associated with *Fst*, *Irx3*, *Klf7*, *Lgr5*, *Tcf4*, and *Tcf12* (**Data S7**). Collectively, our analysis identified LEF1 and MSX1 as transcription factors that could be involved in early pre-granulosa cell differentiation at E11.5. As the supporting cell differentiation progresses at E12.5, a network of transcription regulators become more active in pre-granulosa cells, which undergo significant chromatin remodeling at this time.

### Female-biased *Lef1* and *Msx1* regulation during pre-granulosa differentiation

As LEF1 and MSX1 were identified as potential TFs underlying initial pre-granulosa differentiation, we investigated how *Lef1* and *Msx1* were regulated at the transcript level. We first performed RNA *in situ* hybridization (RNAscope) combined with immunofluorescence of known markers to characterize the cellular localization of *Lef1* and *Msx1* in E11.5 and E12.5 gonads. At E11.5, we observed clear sex differences in *Lef1* and *Msx1* expression. *Lef1* transcripts were abundant in nearly all cell types in the XX gonad, whereas in the XY gonad, *Lef1* was scarcely detected (**Figure 6A, i, ii, v, vi**). Similarly, *Msx1* transcripts were present in most cells in the XX gonad except for the coelomic epithelium, while in the XY gonad, *Msx1* expression was minimal (**Figure 6A, ix, x, xiii, xiv**). At E12.5, *Lef1* remained broadly expressed in almost all cell types in the XX gonad whereas in the XY gonad, *Lef1* expression was restricted to germ cells and interstitial cells (AMH negative/COUP-TFII positive cells) (**Figure 6A, iii, iv, vii, viii**). Similarly, *Msx1* showed a broad expression pattern at E12.5 in an XX gonad, while in the XY gonad *Msx1* expression was low and limited to non-Sertoli cells (**Figure 6A, xi, xii, xv, xvi**).

**Figure 6:**
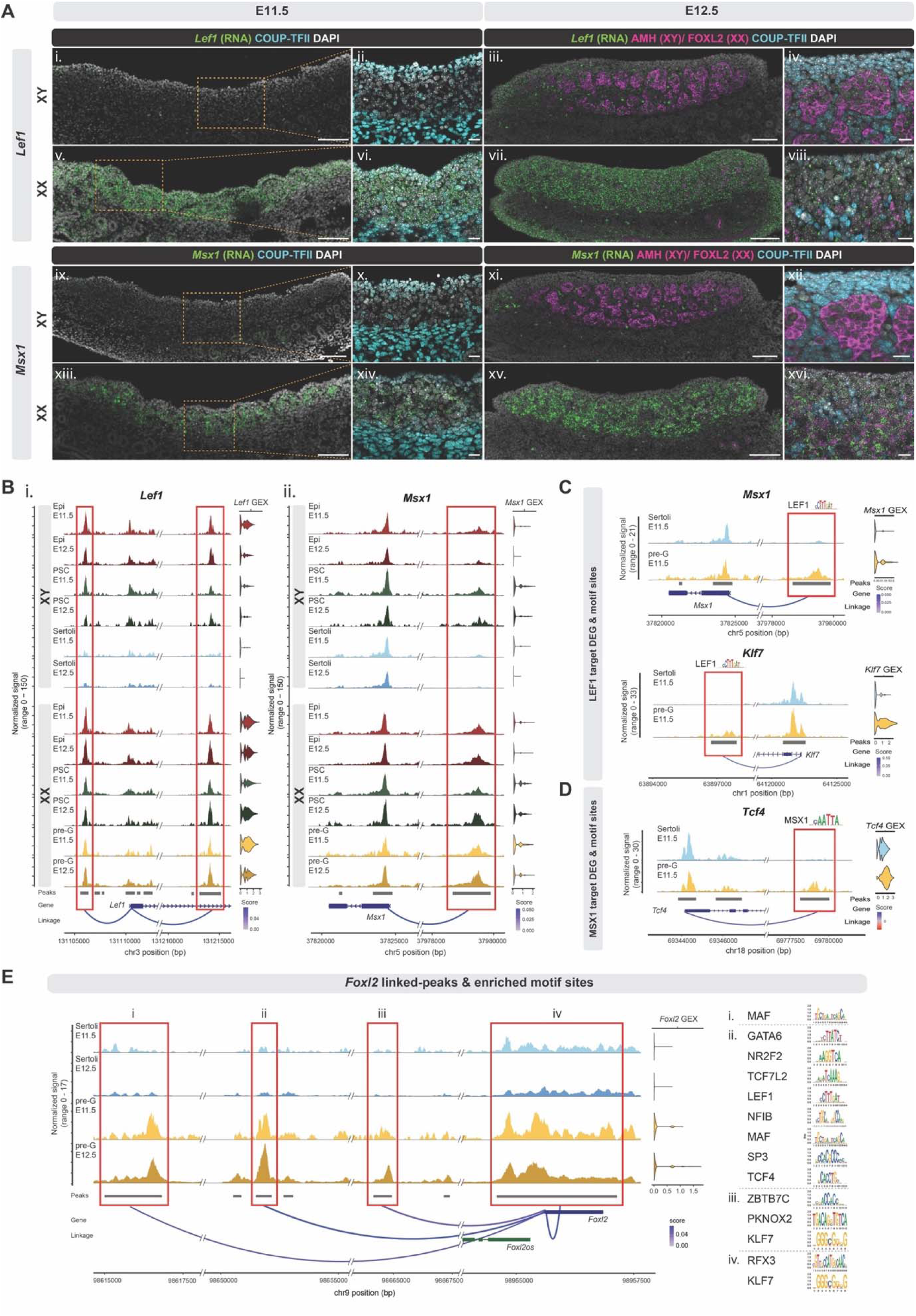
Pre-granulosa cell-biased *Lef1* and *Msx1* expression and regulation. (A) RNA *in situ* hybridization (RNAscope) of *Lef1* and *Msx1* (green) combined with immunofluorescence staining of COUP-TFII (cyan, ii, iv, vi, viii, x, xii, xiv, xvi), AMH (magenta, iii, iv, xi, xii), and FOXL2 (magenta, vii, viii, xv, xvi) on E11.5 and E12.5 XX and XY gonadal sections. Scale bar= 100 mm (10x objective, i, iii, v, vii, ix, xi, xiii, xv) or 20 mm (20x objective, ii, iv, vi, viii, x, xii, xiv, xvi). (B) Peak-gene linkage plots of *Lef1* (i) and *Msx1* (ii) in E11.5 and E12.5 epithelial (EPI), pre-supporting (PSC), Sertoli, and pre-granulosa (pre-G) cells. The red boxes denote called peaks (grey bars) that are linked to gene expression (GEX). Linkage score represents the level of association between chromatin accessibility and gene expression. (C) Peak-gene linkage plots of *Msx1* (i) and *Klf7* (ii) in E11.5 Sertoli and pre-granulosa cells, with LEF1 motif identified within the linked pre-granulosa DAPs (red boxes). (D) Peak-gene linkage plot of *TCF4* in E11.5 Sertoli and pre-granulosa cells, with MSX1 motif identified within the linked pre-granulosa DAPs (red box). (E) Peak-gene linkage plots of *Foxl2* in E11.5 and E12.5 Sertoli and pre-granulosa cells, with identified motifs enriched within each linked peaks (red boxes, i-iv) shown in the right side.

To understand how the sex-dimorphic pattern of *Lef1* and *Msx1* expression is established, we investigated the chromatin regions that were associated with *Lef1* and *Msx1* expression (**Figure 6B**). In the *Lef1* locus, two chromatin regions, one upstream of the *Lef1* promoter and one intragenic, were associated with *Lef1* expression (**Figure 6B, i**). In the XX gonads, these two regions remained accessible in epithelial, pre-supporting, and pre-granulosa cells. However, in the XY gonads, their accessibility was reduced in E12.5 pre-supporting cells and in both E11.5 and E12.5 Sertoli cells (**Figure 6B, i**). For *Msx1*, we identified one chromatin region upstream of the *Msx1* promoter that was associated with its expression (**Figure 6B, ii**). This region also remained accessible in XX epithelial, pre-supporting, and pre-granulosa cells but exhibited reduced accessibility in Sertoli cells (**Figure 6B, ii**). The decreased accessibility among these chromatin regions may explain the reduced expression of *Lef1* and *Msx1* in Sertoli cells. Finally, to explore how granulosa-specific genes including *Foxl2* and *Fst* are regulated, we examined the presence of motifs of all potential regulators identified in **Figure 5E** within accessible peaks associated with these genes. We first identified a LEF1 motif within accessible regions upstream of the *Msx1* promoter and downstream of *Klf7*. These regions were more accessible in pre-granulosa cells than in Sertoli cells (**Figure 6C**). Similarly, an MSX1 motif was identified downstream of *Tcf4* within a chromatin region more accessible in pre-granulosa cells than Sertoli cells (**Figure 6D**). Both KLF7 and TCF4 were among the downstream, secondary factors enriched within E12.5 pre-granulosa “DEG with linked DAPs” (**Figure 5E**). Binding motifs of KLF7 and TCF4, along with other transcription factor such as MAF, GATA6, ZBTB7C, and SP3, were mapped to the accessible peaks associated with *Foxl2* and *Fst* expression (**Figure 6E, S6A**). For *Foxl2*, four chromatin regions were associated with its expression: the *Foxl2* promoter and three upstream regions (**Figure 6E**). Within these regions, the MAF motif was mapped to the first peak (**Figure 6E, i**); motifs such as GATA6, LEF1, SP3, and TCF4 were mapped to the second peak (**Figure 6E, ii**); ZBTB7C and KLF7 were among the motifs identified in the third peak (**Figure 6E, iii**); and RFX3 and KLF7 were mapped to the promoter peak of *Foxl2* (**Figure 6E, iv**). For *Fst*, motifs including TCF4/12, KLF7, ZBTB7C, GATA6, and RUNX1 were found within the three linked peaks closest to its promoter that were associated with *Fst* expression (**Figure S6A**). We also analyzed the chromatin regions linked to the expression of pro-ovarian factors *Wnt4* and *Rspo1*. Enrichment of factors such as MEIS1/2, ZBTB7C, SP3, TCF12, NFIB, and KLF7 was observed within peaks linked to *Wnt4* and *Rspo1* (**Figure S6B, C**). These factors, expressed also in XY precursor cell types, may regulate *Wnt4* and *Rspo1* expression in XY precursor cells before the linked chromatin regions become inaccessible in Sertoli cells (**Figure S6B, C**). Unlike *Foxl2* and *Fst*, *Wnt4* and *Rspo1* were expressed in both XX and XY precursor cell types (**Figure S6B, C**). *Wnt4* was differentially expressed between E11.5 XX and XY coelomic epithelial cells; however, its expression was low in the epithelium (**Data S3**). *Wnt4* was upregulated in pre-supporting cells of both sexes, further increasing in pre-granulosa cells while decreasing in Sertoli cells. Meanwhile, *Rspo1* was present in E11.5 XX and XY epithelial cells and pre-supporting cells. Its expression was upregulated in pre-granulosa cells but downregulated in Sertoli cells, becoming a DEG only at this stage (**Data S3**). Together, these findings suggest a potential mechanism by which LEF1 and MSX1 may initiate a transcription network cascade, eventually leading to upregulation of downstream secondary factors that collectively facilitate pre-granulosa gene expression and differentiation.

## Discussion

Our understanding of the molecular events underlying sex determination- how the testis or ovary pathways are initiated from the bipotential state- has greatly advanced thanks to high-throughput technologies. These approaches enable the discovery of changes in gene expression in individual cells or cell types across different developmental timepoints (27, 31, 36-38), as well as the association between chromatin states and gene expression in bulk samples collected before and after sex determination (23-25). Nevertheless, simultaneous analysis of gene expression and chromatin status in various cell types in the gonads remains a challenge. To investigate the transcriptional regulatory network driving the fate establishment of supporting cells, we employed single-nucleus joint transcriptomics and chromatin accessibility assays. This approach allows the identification of active transcription factor motifs through linking accessible chromatin regions to gene expression within the same cells. Moreover, its single-cell resolution captures the transition from precursor cells to early differentiated supporting cells at the same timepoints (e.g. epithelial, pre-supporting, Sertoli, and pre-granulosa cells at E11.5, the beginning of sex determination). Integrating these data, we explored the transcriptional networks of supporting cell differentiation by comparing across sexes and cell types. Our analyses revealed a strong correlation between chromatin modulation and gonadal cell differentiation. Specifically, Sertoli and pre-granulosa cells acquired sex-specific chromatin regions during sex determination, accompanied by the deposition of active histone marks. Sertoli cells exhibited an active increase in the accessibility of chromatin regions linked to Sertoli cell-enriched genes, while reducing accessibility in chromatin regions associated with pre-granulosa-specific genes, creating a chromatin state that is markedly distinct from other supporting cells. In contrast, less pronounced changes were observed in pre-granulosa cells, which shared a more similar chromatin profile with pre-supporting cells. Notably, active chromatin remodeling occurred between E11.5 and E12.5 in both cell types, with a marked increase in the proportion of differential chromatin regions corresponding to sex-specific gene expression. Based on these changes, we explored the transcriptional regulatory networks underlying the differentiation of both Sertoli and pre-granulosa cells.

### Validity of single-nucleus multiomics in cell type identification and peak-gene association

Our single-nucleus multiomics data identified all gonadal cell types with cell composition that aligns with published single-cell RNA-seq datasets (27, 31). Compared to single “cell” RNA-seq, expression of certain genes (e.g., *Foxl2* in pre-granulosa cells) appeared lower in our single “nucleus” RNA-seq data, likely because only nuclear mRNA was captured using the multiome assay. Nonetheless, we suggest that nuclear mRNA may reflect active transcription more closely compared to total mRNA analyzed with single-cell RNA-seq, as nuclear mRNA represents higher proportion of newly transcribed genes (39). By correlating accessible chromatin regions with gene expression, we captured dynamic changes in chromatin peaks that explain different gene expression patterns across cell types. Using the canonical pre-granulosa marker *Fst* and Sertoli marker *Amh* as examples, we observed distinct patterns in how female and male-specific chromatin peaks are established: chromatin accessibility at *Fst* regions gradually increased across epithelial, pre-supporting, and pre-granulosa cells but remained inaccessible in Sertoli cells. In contrast, the *Amh* regions were inaccessible in precursor cells and then became abruptly accessible in Sertoli cells. These findings highlight the continuous and gradual differentiation of pre-granulosa cells, versus a rapid transition in Sertoli cell differentiation. In setting the parameters for linkage analysis, we noticed that the analysis is sensitive to the composition of the dataset. We obtained the most peak-gene linkages when the analysis was performed on the entire dataset, encompassing all cell types, compared to performing the analysis within individual cell types. Our approach allowed for the identification of all potential regulatory chromatin regions not limited to supporting cells. Nonetheless, the linkage results may differ if the analysis was restricted to Sertoli and pre-granulosa cells alone.

### Sex- and cell type-specific chromatin remodeling during gonadal differentiation

Changes in chromatin accessibility are associated with supporting cell differentiation, as demonstrated by ATAC-seq performed on FACS-sorted gonadal somatic cells at timepoints before and after sex determination (23, 25). In our analysis, we observed that chromatin states correlated with cell type-specific gene expression among major cell types, suggesting the involvement of chromatin remodeling in not only Sertoli and pre-supporting cell differentiation, but also in the emergence of supporting-like cells (27) and interstitial cell differentiation. While we observed exceptions to this correlation when comparing directly between RNA and ATAC clusters, we reason that they were due to the resolution of the clustering analysis used. When we explored further into the chromatin states during supporting cell differentiation across timepoints, we noticed that epithelial and pre-supporting cells at E11.5 exhibited minimal sex differences, reflecting the bipotential state of somatic precursor cells. Further, we detected the highest number of differentially accessible peaks (DAPs) between Sertoli and pre-granulosa cells. Specifically, Sertoli cells exhibited hundreds of Sertoli-specific chromatin regions readily by E11.5. The acquisition of these Sertoli DAPs was driven by an active increase in chromatin accessibility, particularly in distal and intronic regions, as shown by increased accessibility compared with epithelial and pre-supporting cells. In pre-granulosa cells, the acquisition of DAPs, also in majority in distal and intronic regions, occurred pre-dominantly at E12.5. In contrast, the acquisition of pre-granulosa DAPs was a result from a combination of Sertoli cells decreasing accessibility and pre-granulosa cells gaining accessibility in these regions. These results support the model in which SRY become active in XY supporting cell progenitors around E11.0 (40). The action of SRY, its downstream effector SOX9, and other transcription factors lead to the opening of previously condensed chromatin regions by E11.5. Without SRY and the activation of its downstream pathway, chromatin changes in pre-granulosa cells were not observed until later developmental timepoint, potentially facilitated through other chromatin remodelers unique to the XX environment. By analyzing histone modifications marking active enhancers and promoters, we observed a drastic increase in H3K4me3 and H3K27ac marks within both Sertoli and pre-granulosa DAPs when comparing E10.5 FACS-sorted pre-supporting cells to E13.5 Sertoli and pre-granulosa cells (25). These findings further support that Sertoli and pre-granulosa-specific DAPs identified in our dataset may be accompanied by active transcription. While our analysis suggests that Sertoli and pre-granulosa cells acquire these active histone marks within either E11.5 or E13.5 DAP regions by E13.5, whether the changes in histone modifications occur readily by E11.5 will need to be determined though histone profiling of cells at E11.5. In another words, whether the deposition of histone modifications drives the acquisition of DAPs and the expression of Sertoli and pre-granulosa-enriched genes, or occurs afterwards as a reflection of the transcriptional state, remains to be determined (41).

Sertoli and pre-granulosa-enriched genes are marked initially as bivalent with both active H3K4me3 and repressive H3K27me3 histone modifications at their promoter regions, and this bivalency is lost after sex determination (25). In our analysis, however, we did not observe significant changes in the repressive H3K27me3 modifications within Sertoli and pre-granulosa DAPs. We reason that this was due to the majority of DAPs being located within distal intergenic and intragenic regions. Moreover, we found that known bivalent promoters of genes including *Sox9*, *Dmrt1*, and *Fgf9*, were not DAPs between Sertoli and pre-granulosa cells and therefore were not included in the analysis. These results provide a glimpse into the dynamic changes in distal regulatory regions, in addition to the action of bivalent promoters, occurring during Sertoli and pre-granulosa cell differentiation.

### Sertoli and pre-granulosa cells exhibit increased DEG associated with open chromatin during E11.5 to E12.5 transition

To understand how sex-specific genes are established over time, we analyzed DEGs and their association with open chromatin regions between sexes across timepoint. We categorized DEGs based on whether they are associated with any chromatin accessible peaks, and if so, whether this peak is specific to one sex in that cell type. We observed that Sertoli cells and pre-granulosa cells exhibited the highest level of “DEG with linked DAP”, reflecting how Sertoli and pre-granulosa cells possessed the highest number of DAPs. While the level of increase in DAP in Sertoli and pre-granulosa cells from E11.5 to E12.5 does not correlate with changes in the number of DEG, we propose that this may reflect the ability of multiple DAPs to function as regulatory elements for a single gene. “DEG with linked DAP” was also detected in epithelial and mesenchymal cells, consistent with the levels of DAPs observed in these cell types. Collectively, these findings suggest that some sex-specific DEGs of Sertoli, pre-granulosa, epithelial, and mesenchymal cells are regulated at the chromatin level. In contrast, the majority of DEGs in pre-supporting and supporting-like cells were linked to non-DAP. This suggests that these genes may be regulated independently of chromatin state change. For example, the linked chromatin region is equally accessible in both sexes, while gene expression is regulated by different sex-specific transcription factors. We also observed that more than 30% of DEGs between Sertoli and pre-granulosa cells were not associated with any peaks at all. As the linkage analysis was performed within a distance of transcription start site ± 500Kb, distal regulatory elements beyond this linkage distance (e.g., Enh13 in Sertoli cells (42)) were not considered in the analysis. Therefore, these genes could either be regulated by sex-specific transcription factors, or through distal enhancers beyond the linkage distance, which could be explored in future studies using chromosome conformation capture-based techniques (43).

Perhaps most strikingly, we found a large proportion of “DEGs with linked non-DAPs" at E11.5 converted to “DEG with linked DAP” at E12.5 in both Sertoli and pre-granulosa cells. This pattern may result from a small subset of cells expressing high level of the gene at E11.5, rendering it a DEG, while the chromatin accessible peaks of these genes in these cells do not reach statistical significance when analyzed in combination with the rest of the cells as a group. By E12.5, most cells increase the accessibility of the associated chromatin regions, or decrease it in the opposite sex, establishing it as a DAP. Alternatively, the DEGs with linked non-DAP at E11.5 may be regulated through mechanisms independent of chromatin accessibility. In either scenario, these dynamic changes in the association between chromatin accessibility and gene expression allowed us to identify computationally the enriched transcription factors underlying this active transition.

### Identification of known and new regulators of Sertoli cell differentiation

At E11.5, when sex determination begins, motif scan and linkage analysis revealed that SOX motifs were enriched in Sertoli DAPs linked to *Dmrt1* expression, suggesting potential direct regulation of *Dmrt1* through SOX factors at the beginning of Sertoli differentiation. This result aligns with existing knowledge that SOX9 and DMRT1 play critical roles in Sertoli cell specification and fate maintenance (44, 45). Notably, despite the similarity between SOX and DMRT1 motifs, clustering based on downstream linked genes indicated that these two genes regulate different target genes, also consistent with our understanding of their distinct roles in Sertoli cell function.

At E12.5 when the supporting cell lineages are established, we identified other transcription factors such as NR6A1, thyroid hormone receptor alpha (THRA), JUN, and IRF2 with high predicted motif activity. NR6A1 shares a similar motif with NR5A1 but exhibited higher gene expression that *Nr51a* in Sertoli cells, particularly during the E11.5 pre-supporting to Sertoli transition. Also known as the Germ Cell Nuclear Factor (GCNF), NR6A1 is highly expressed in germ cells (26) and is implicated in regulating spermatogenesis and oogenesis (46, 47). Its enrichment and differential expression in Sertoli cells may indicate an additional and yet-to-be-identified role in mediating Sertoli differentiation. Our analysis further suggested that THRA, JUN, and IRF2 (48) may regulate early Sertoli differentiation. Sertoli cells express thyroid hormone receptors and respond to thyroid hormone signaling, which drives their proliferate during the neonatal period (49). JUN is also expressed in Sertoli cells and may interact with PA1 to regulate junctional protein expression (50). Although IRF2 is known for its role in immune regulation and cell growth (51, 52), its specific involvement in Sertoli cell differentiation has yet to be defined.

### Joint transcriptomic and chromatin accessibility assays identified transcriptional regulatory networks of pre-granulosa cell differentiation

We identified LEF1 and MSX1 that could function as upstream regulators of pre-granulosa cell differentiation. LEF1 is a member of T cell factor/ lymphoid enhancer factor (TCF/LEF) family and acts as a key transcription factor downstream of the WNT signaling pathway (53). The pathway, specifically its factors WNT4 and RSPO1, play essential roles in early recruitment of pre-supporting cells from the epithelium in both sexes in addition to regulating ovarian differentiation (54). The fact that the LEF1 motif was already active in epithelial cells, preceding the upregulation of *Wnt4* in pre-supporting cells, suggested that it could be regulated by a different WNT factor, such as *Wnt5a* (38). Alternatively, LEF1 could be regulated through other pathways or independently (55, 56). The exact mechanism of how *Lef1* is regulated in precursor cell types require further characterization. The LEF1 motif remained active in XX pre-supporting and pre-granulosa cells, while its activity was downregulated in XY cells. As a member of the LEF/TCF family, LEF1 shares similar motifs with TCF7L1 and TCF7L2, both of which were identified as potential secondary factors in pre-granulosa differentiation.

Other than transmitting WNT signaling, a potential action of LEF1 is to directly modulate *Msx1* expression (57). *Msx1* encodes a homeodomain transcription factor involved in meiosis initiation (58). Double knockout of *Msx1* and *Msx2* exhibited reduced meiotic germ cells in mouse ovaries (58). However, its role in somatic cell differentiation in the ovary remains unknown. MSX1 shares similar motif with LHX9 and EMX2, both of which are involved in coelomic epithelial proliferation and early fate establishment (59, 60), and were identified among the secondary factors in our analyses. The expression of *Lhx9* and *Emx2* persisted in XX pre-supporting cells and pre-granulosa cells but was downregulated in XY cells.

Among the secondary factors, we identified including RUNX1, GATA6, TRPS1, and NF1A/B as transcription factors with high predicted motif activity in pre-granulosa cells. RUNX1 marks pre-supporting cells in both sexes, and then becomes differentially expressed in pre-granulosa cells (16). Along with FOXL2, RUNX1 maintain pre-granulosa identity (16). GATA6, a member of the GATA transcription factor family, interact with GATA4 in regulating early folliculogenesis in mice (61). While GATA6 and GATA4 share similar motif, GATA4 was differentially expressed in Sertoli cells and was not identified as an enriched motif in pre-granulosa cells. Nevertheless, GATA4 was expressed and predicted to have high motif activity in pre-granulosa cells. TRPS1, another member of the GATA transcription factor family, also shares a similar motif with GATA6 and GATA4. While TRPS1 has been associated with high-grade serous ovarian carcinoma (62), its role in early ovarian development remains underexplored. Similarly, members of the Nuclear Factor I family, including NFIA and NFIB were identified in our analyses as potential secondary regulators. These two factors were implicated in ovarian cancer metastasis and drug resistance (63, 64); however, their roles in ovarian differentiation have not been defined. It is worth noting that none of these secondary factors in pre-granulosa cells were negatively-linked to genes encoding the identified Sertoli TFs. Whereas several Sertoli TF motifs were enriched within peaks negatively-linked to *Bnc2*, *Zbtb7c*, *Pbx1*, *Tcf4*, and *Gli3*. These observations suggest that Sertoli cells actively suppress pre-granulosa fate through negatively regulating the expression of pre-granulosa-enriched transcription factors, but not vice versa.

The joint single-nucleus RNA and ATAC multiomics leverage both open chromatin patterns and gene expression data, enabling the identification of putative transcription factors binding to open chromatin regions. Therefore, the factors proposed in this study were limited to transcription factors identified through motif analysis within DAPs between Sertoli and pre-granulosa cells. Other factors involved in signaling transduction, post-translational modifications, or metabolic regulation that are not necessarily transcription factors were not included in the study. The -KTS isoform of *Wt1* was reported to be essential for pre-granulosa cell differentiation (17). Without total RNA or long-read sequencing, we were not able to capture and quantify the variant in our dataset. Moreover, *Wt1* was not a DEG in supporting cells between sexes across timepoints. Finally, future analyses could explore the role of molecules secreted by supporting cells in activating pathways that regulate target gene expression in other cell types.

Collectively, we present a robust dataset and downstream analysis that enables the interrogation of chromatin states and gene expression at single-cell resolution during gonadal differentiation. By comparing the transcriptional regulatory network between Sertoli and pre-granulosa cells, we propose a model for the less well-defined regulation of pre-granulosa genes. This model involves how LEF1, a downstream effector of the WNT signaling pathway, works in conjunction with its potential target, MSX1, as upstream regulators of pre-granulosa differentiation. LEF1 and MSX1 may directly regulate the expression of secondary factors to promote the expression of key pre-granulosa genes such as *Foxl2* and *Fst*. This dataset, along with the putative mechanisms and factors identified through computational analyses, is particularly valuable for studying transcriptional regulatory networks underlying gonadal cell fate transitions, including the differentiation of Sertoli, pre-granulosa, supporting-like cells, and the establishment of interstitial lineage, advancing our understanding of the various causes of human DSD and other reproductive system disorders.

## Materials and Methods

### Animals

Twelve- to Sixteen-week-old female mice bred from homozygous Rosa-tdTomato9 females (JAX 007909) (B6.Cg-Gt(ROSA)26Sor<tm9(CAG-tdTomato)Hze>/J) crossed to homozygous Nr5a1-cre males (B6D2-Tg(Nr5a1-cre)2Klp, provided by the late Dr. Keith Parker (65)) were used for the 10x Multiome assay. C57BL/6J (JAX 000664) female mice of the same age were used for RNA *in situ* hybridization. The day of detection of vaginal plug after time mating was considered embryonic day (E) 0.5. Food (NIH-31M, Harlan Teklad) and water were given *ad libitum* and the mice were kept in a 12-h light, 12-h dark cycle with temperature ranging 21–23 °C and relative humidity ranging 40 to 50%. All animal studies were conducted in accordance with the NIH Guide for the Care and Use of Laboratory Animals and approved by the National Institute of Environmental Health Sciences (NIEHS) Animal Care and Use Committee.

### Gonadal sample collection and preparation

Whole fetal mouse gonads were collected at E11.5, E12.5, and E13.5 in ice-cold phosphate-buffered saline (PBS) using insulin needles under a fluorescence microscope (Leica M165 FC). As tdTomato fluorescence is only present in the gonad but not the mesonephros, it was used to facilitate the dissection of the gonad away from the mesonephros. Tail somite counting was performed on E11.5 embryos which ranged from 17 to 23 somites. The sexes of E11.5 and E12.5 embryos were determined using amnion staining (66) and confirmed with PCR using primers that recognize the Y chromosome. Gonads of the same sex and age were pooled in PBS with 0.4% bovine serum albumin (BSA, Signa-Aldrich) and immediately frozen in liquid nitrogen until nuclei isolation. Nuclei isolation was performed using the 10X Genomics Chromium Nuclei Isolation Kit (Cat# 1000494) following the manufacturer’s protocol. The total numbers of gonads used per sex and age are listed in **Table S2**.

### Library preparation for single-nucleus RNA-sequencing and ATAC-sequencing

The 10X Genomics Chromium Next GEM Single Cell Multiome ATAC + Gene Expression Library Preparation Kit (Cat# 1000284) was used to generate a 10X barcoded library of mRNA and transposed DNA from individual nuclei. Libraries were prepared from 3-19 biological replicates (pairs of gonads) and two technical replicates per embryonic stage and sex (**Table S2, Figure 1A, B**), averaging 8,687 nuclei per sample for sequencing with Illumina NOVAseq to a minimum sequencing depth of 129,120,442 raw reads and 22,406 read pairs/nucleus (**Table S2, Figure S1A, B**).

### Data preprocessing for single-nucleus RNA-seq and ATAC-seq libraries

*CellRanger-ARC* (v3.0) (67, 68) was used for count, alignment to mm10 reference genome, filtering, cell barcode and UMI counting (**Table S2, Figure S1C**). Barcode swapping correction was performed for all libraries (69). FASTQ files of the corrected cell count matrices were generated and the data were analyzed using *Seurat* (v. 4.3.0) (70) and *Signac* (v. 1.9.0) (71) in R (v. 4.2.1) with default settings unless otherwise noted. For snRNA-seq preprocessing, doublets within each dataset were removed with the *scDblFinder* (v. 1.10.0) (72) package using default methods. Ambient RNA was removed using the *celda/decontX* (v. 1.12.0) (73) package, with the maximum iterations of the EM algorithm set at 100. Datasets of gonadal cell populations from each embryonic stage and sex were combined into one *Seurat* object using the “merge” function. As batch effect was not observed in our dataset, we did not perform further integration or batch correction (**Figure S1D**). A subset of the data was generated using the following cutoffs: nCount_RNA > 1000 & nCount_RNA < 25,000 & percent.mt < 25. The data was normalized using the “SCTransform” function in *Seurat*. For snATAC-seq data preprocessing, peak calls from *CellRanger* were used to generate one combined peak set using the function “reduce” and a subset of the data was generated using the following cutoffs: peak size > 20 bp & peak size < 10,000 bp. Fragment objects were created using the *Signac* package and combined with the *Seurat* object. A subset of the snATAC-seq data was generated using the following cutoffs: nCount_ATAC < 100,000 & nCount_ATAC > 1000 & nucleosome_signal < 2 & TSS.enrichment > 1.

### Computational sex filtering

We applied computational filters to confirm the sex of the cells in each dataset based on Y-linked gene expression and fragments mapped to the Y chromosome (chrY). Y-linked genes that were expressed in our dataset including *Kdm5d, Eif2s3y, Uty,* and *Ddx3y,* were used to create a module score using the *UCell* package (v. 2.0.1) (74). XY cut-off value was set at one standard deviation below the mean module score of all XY E13.5 samples. Whereas XX cut-off value was set at one standard deviation above the mean module score of all XX E13.5 samples. Next, we examined the peaks called on chrY outside of the pseudoautosomal region. In total, 21 peaks were identified between chrY 1-90,000,000 bp. E11.5 and E12.5 samples with 0 fragment counts within our defined chrY peak region and Y-linked gene expression module score < 1 standard deviation were removed from the “XY” datasets. E11.5 and E12.5 samples with > 0 fragment counts within our defined chrY peak region or Y-linked gene expression module score > 1 standard deviation were removed from the “XX” datasets.

### Clustering and cell type annotation

We performed linear dimensional reduction using the “RunPCA” function and non-linear dimensional reduction using the “RunUMAP” function with dims = 1:15 on the scaled snRNA-seq data. Cell clustering was then performed using the “FindNeighbors” function with dims= 1:15 and the “FindClusters” function with resolution= 0.3. This generated in total 14 cell clusters. We combined established marker gene expression, sex and age composition for cell type annotation. Specifically, primordial germ cell (PGC) cluster is positive of *Ddx4* and *Pou5f1* expression; pre-granulosa (pre-G) is an XX cluster and is positive of *Fst* and *Foxl2* (15); Sertoli is an XY only cluster and is positive of *Amh* and *Sox9* (30); pre-supporting cell (PSC) shares similar gene expression with pre-G and Sertoli including *Wnt4*, *Wnt6*, *Gata4*, and *Wt1*, and is enriched in both sexes at E11.5 (16); supporting-like cells (SLC) is positive of *Pax8* and *Slc25a21* (27); epithelial cluster is positive of *Upk3b* and *Aldh1a*2 (28, 29); Leydig cluster is positive of *Cyp11a1* and *Hsd3b1* and comprised of only XY samples; mesenchymal (Me) cluster is positive of *Ptn* and *Nr2f2*, while the expression pattern is similar to interstitial progenitor (IP) cluster, IP is more closely related to the Leydig cluster; erythrocyte cluster is positive of *Hba-a1*; immune cluster is positive of *Cx3cr1* and *Fcgr1*; endothelial cluster is positive of *Pecam1* but negative of PGC gene expression; mesonephric tubule (MT) is positive of *Pax2*. (2, 27, 31, 37)

### Peak calling by cluster and linkage analysis

We performed peak calling using MACS2 (v. 2.2.7.1) (32) on individual RNA clusters to obtain cluster-specific peaks. Peaks on nonstandard chromosomes were removed using the “keepStandardChromosomes” function in *Signac* with the option ‘pruning.mode = “coarse”’ and peaks in mm10 genomic blacklist regions were removed. Counts in each peak were determined using *Signac* functionalities. The data was normalized by latent semantic indexing (LSI) using the “RunTFIDF”, “FindTopFeatures”, and “RunSVD” functions in *Signac*. Visualization was performed using UMAP with dims = 2:30, reduction = ‘lsi’, and algorithm = 3. MACS2 called peaks were linked to scaled gene expression data using the entire dataset with the “LinkPeaks” function in *Signac* for quantifying the correlation between peak accessibility with gene expression.

### Identification of differentially expressed genes and differentially accessible peaks

Differentially expressed genes (DEG) in each cluster were determined using the “FindAllMarkers” function in *Seurat* based on the Wilcoxon Rank Sum test with min.pct = 0.25 and logfc.threshold = 0.25. DEGs between male and female somatic clusters at each stage were identified with the “FindMarkers” function in *Seurat* with min.pct = 0.25 and logfc.threshold = 0.25. Differentially accessible peaks (DAP) in each cluster were determined using the “FindAllMarkers” function based on the LR test with min.pct = 0.05 and logfc.threshold = 0.25. DAPs between male and female somatic clusters at each timepoint were identified with the “FindMarkers” function with min.pct = 0.01 and logfc.threshold = 0.1. To annotate linkage between DEGs and DAPs, when at least one DAP is significantly (p < 0.05) linked to a DEG, the DEG is annotated “DEG with linked DAP”; when any of the significantly linked peak to a DEG is not a DAP, the DEG is annotated “DEG with linked non-DAP”; when a DEG has no significantly linked peak it’s annotated “DEG with no linked-peak” (**Data S3**). In case of DAP, when at least one DEG is significantly (p < 0.05) linked to a DAP, the DAP is annotated “DAP with linked DEG”; when any of the significantly linked gene to a DAP is not a DEG, the DAP is annotated “DAP with linked non-DEG”; when a DAP has no significantly linked gene it’s annotated “DAP with no linked-gene” (**Data S2**).

### Peak annotation and histone mark analysis

Peak annotation was performed using ChIPseeker (v. 1.42.0) (75, 76). To analyze chromatin accessibility within cell type-specific DAPs, function “CountsInRegion” in *Signac* was applied. To examine histone marks overlapping cell type-specific DAPs, we applied published ChIP-seq datasets of H3K27ac (GSE118755) (23), H3K4me3, and H3K27me3 (GSE130749) (25) on FACS-sorted gonadal cells. Genome coordinates were converted from mm9 to mm10 assembly using the UCSC Genome Browser (77) LiftOver tool before the analysis.

### Transcription factor binding site motif enrichment analysis

DNA sequence motif analysis of DAPs linked to DEGs was performed using the “FindMotifs” function in *Signac*. Motif position frequency matrices (PFM) for vertebrates were obtained from the 2022 JASPAR Core database (78) with in total 1956 elements. Background peaks were selected to match the GC content in the peak set by using the “AccessiblePeaks,” “GetAssayData”, and “MatchRegionStats” functions in *Signac*. Enriched motifs were filtered with p.adjust < 0.05 and fold.enrichment > 1.25. *MotifScan* (v. 1.3.0) (34) was used to determine the genomic position of linked peaks to DEGs to identify predicted target genes of transcription factors.

### RNA *in situ* hybridization and immunostaining

Gonads from E11.5 and E12.5 C57BL/6J mice were dissected and fixed in 4% paraformaldehyde overnight at 4 °C before being processed for paraffin embedding and sectioning into 5 μm slices with standard protocols. Multiplex fluorescent reagent kit v2 (RNAscope™, Advanced Cell Diagnostics) was used for RNA *in situ* hybridization. All procedures were carried out according to manufacturer’s recommendations. Specifically, RNAscope™ probes Mm-Lef1 (Cat# 441861) and Mm-Msx1 (Cat# 421841, Advanced Cell Diagnostics) were applied. Before DAPI staining, the slides were blocked in blocking buffer (5% donkey serum/0.1% Triton X-100 in PBS) for 1 h at room temperature and counterstained with anti-COUP-TFII (1:200, R&D Systems, Cat# PP-H7147-00), anti-AMH (1:500, Santa Cruz, Cat# sc-6886), or anti-FOXL2 (1:200, Novus Biological, Cat# NB100-1277) antibodies diluted in blocking buffer overnight at 4 °C. The next day, the slides were washed and incubated with Donkey anti-mouse (1:200, Invitrogen, Cat# A10037) and Donkey anti-goat (1:200, Invitrogen, Cat# A21447) secondary antibodies diluted in blocking buffer incubation for 1 h at room temperature. The slides were imaged with a Zeiss confocal microscope (Zeiss).

### Statistical analyses

No sample size calculation was performed. Statistical analyses were considered significant if *p* < 0.05.

## Acknowledgments

We are grateful for the current and past members of the Yao lab, especially B. Nicol, C. Amato, and S. Ratan for their valuable input and assistance with experiments. We thank the NIEHS Comparative Medicine Branch for mouse colony maintenance and to the Epigenomics and DNA Sequencing Core and the Integrative Bioinformatics Support Group, especially S. Grimm, B. Bennett, and F. Day for their assistance with sequencing and data analysis.

## Author contributions

Conceptualization: YYC, HHCY

Methodology: KR, AKA, XX, BP, MAE

Investigation: YYC, AKA

Visualization: YYC

Supervision: HHCY

Writing: YYC, HHCY

## Data and materials availability

Raw sequencing data and code will be deposited upon publication.

## Supplementary Materials

Figures. S1 to S6

Tables S1 to S2

Data S1 to S7

