## Supplementary files and tables for "Single-nucleus multiomics of murine gonadal cells reveals transcriptional regulatory network underlying supporting lineage differentiation"

Supplementary Materials

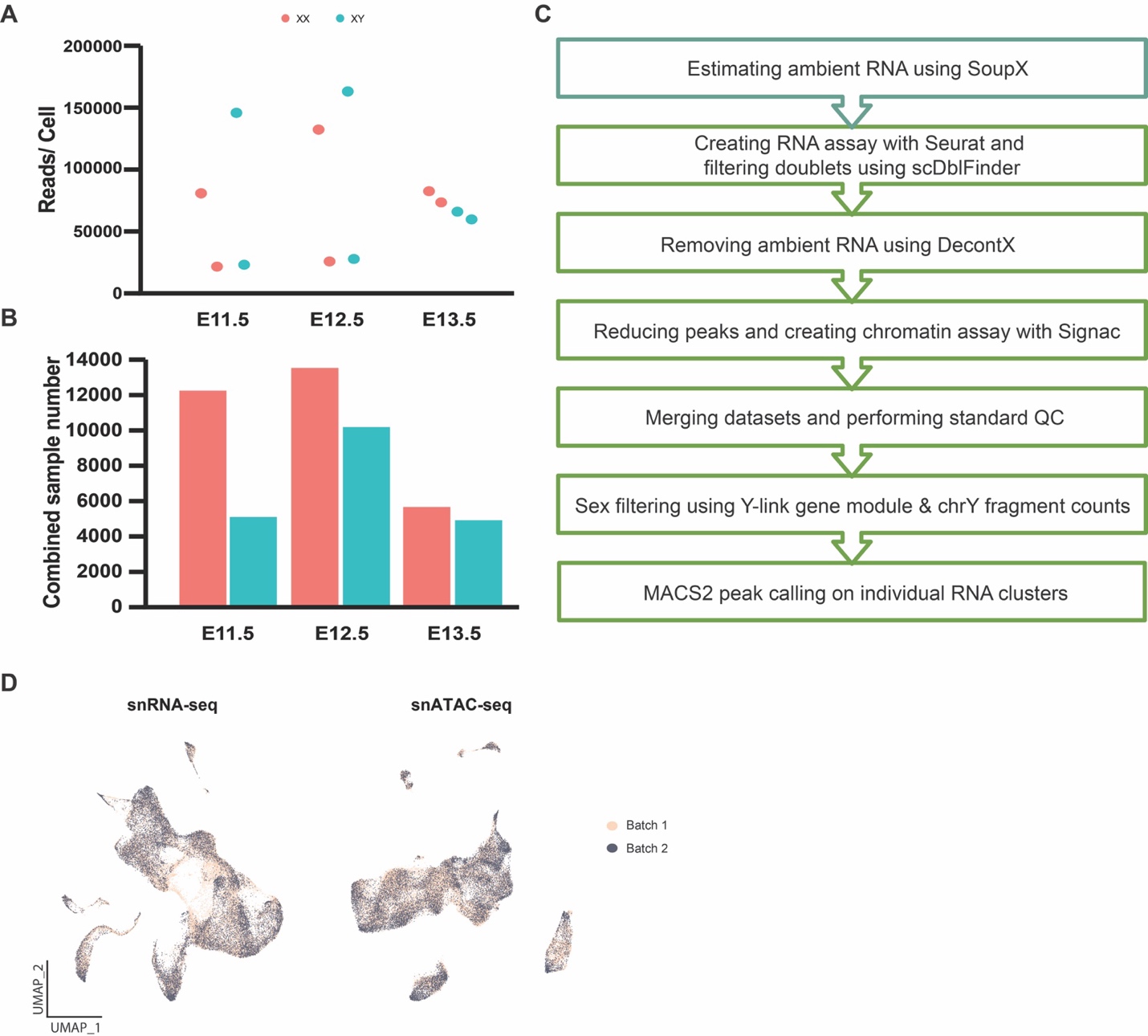

**Figure S1.** **Joint single-nucleus multiomics pre-analysis pipeline.** (A-B) Sequencing depth (A) and nuclei number (B) of individual sample. (C) Pre-analysis pipeline, including ambient RNA removal, doublets filtering, sex filtering, and peak calling. (D) UMAP visualization of snRNA-seq and snATAC-seq data, color-coded by batch.

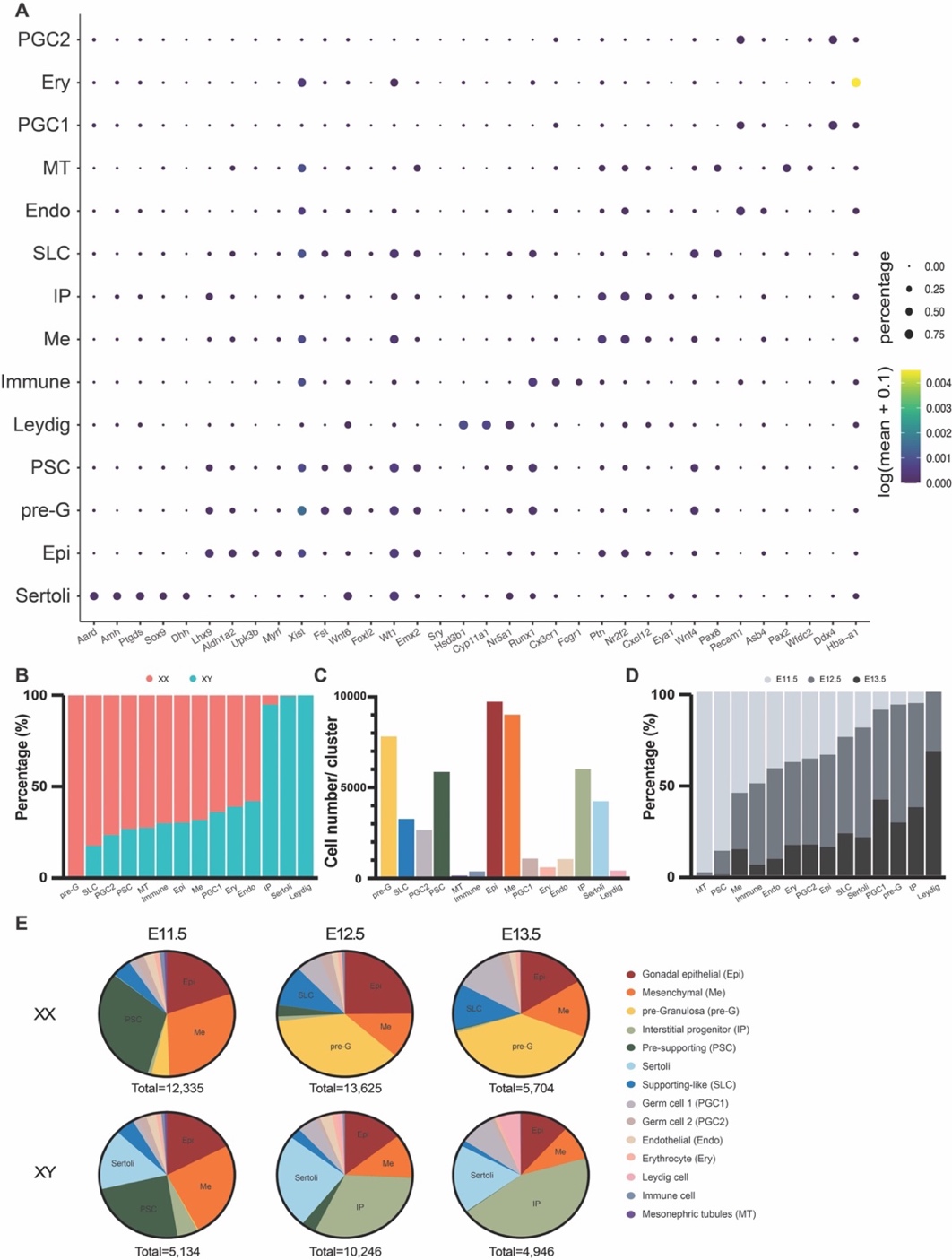

Figure S2. Cell type annotation and distribution. (A) Gene expression profile of the top DEGs identified in each cell type, annotated through RNA clustering. (B) Percentage of cells from each sex within each cell type. (C) Total number of nuclei in each cluster. (D) Percentage of developmental timepoints in each cluster. (E) Percentage of cell types in XX and XY gonads at E11.5, E12.5, and E13.5. Annotations for the clusters are: Gonadal epithelial (Epi), Mesenchymal (Me), Pre-granulosa (pre-G), Interstitial progenitor (IP), Pre-supporting (PSC), Supporting-like cell (SLC), Germ cell (PGC), Endothelial cell (Endo), Erythrocyte (Ery), Mesonephric tubule (MT).

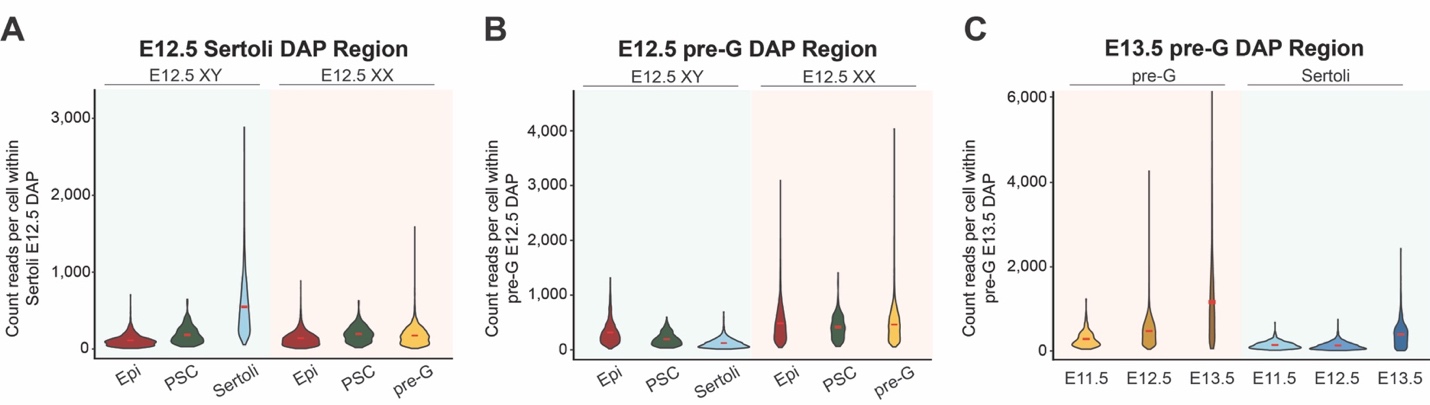

**Figure S3. Chromatin accessibility of Sertoli and pre-granulosa DAPs in precursor and differentiated supporting cells.** (A) Chromatin accessibility, measured by total count reads per cell, within E12.5 Sertoli DAP regions in E12.5 XY and XX epithelium (Epi), pre-supporting cells (PSC), Sertoli, and pre-granulosa cells (pre-G). (B) Total count reads per cell within E12.5 pre-granulosa DAP region in E12.5 XY and XX epithelium, pre-supporting, Sertoli, and pre-granulosa cells. (C) Total count reads per cell within E13.5 pre-granulosa DAP regions (10,124 unique peaks) in pre-supporting and Sertoli cells throughout developmental timepoints.

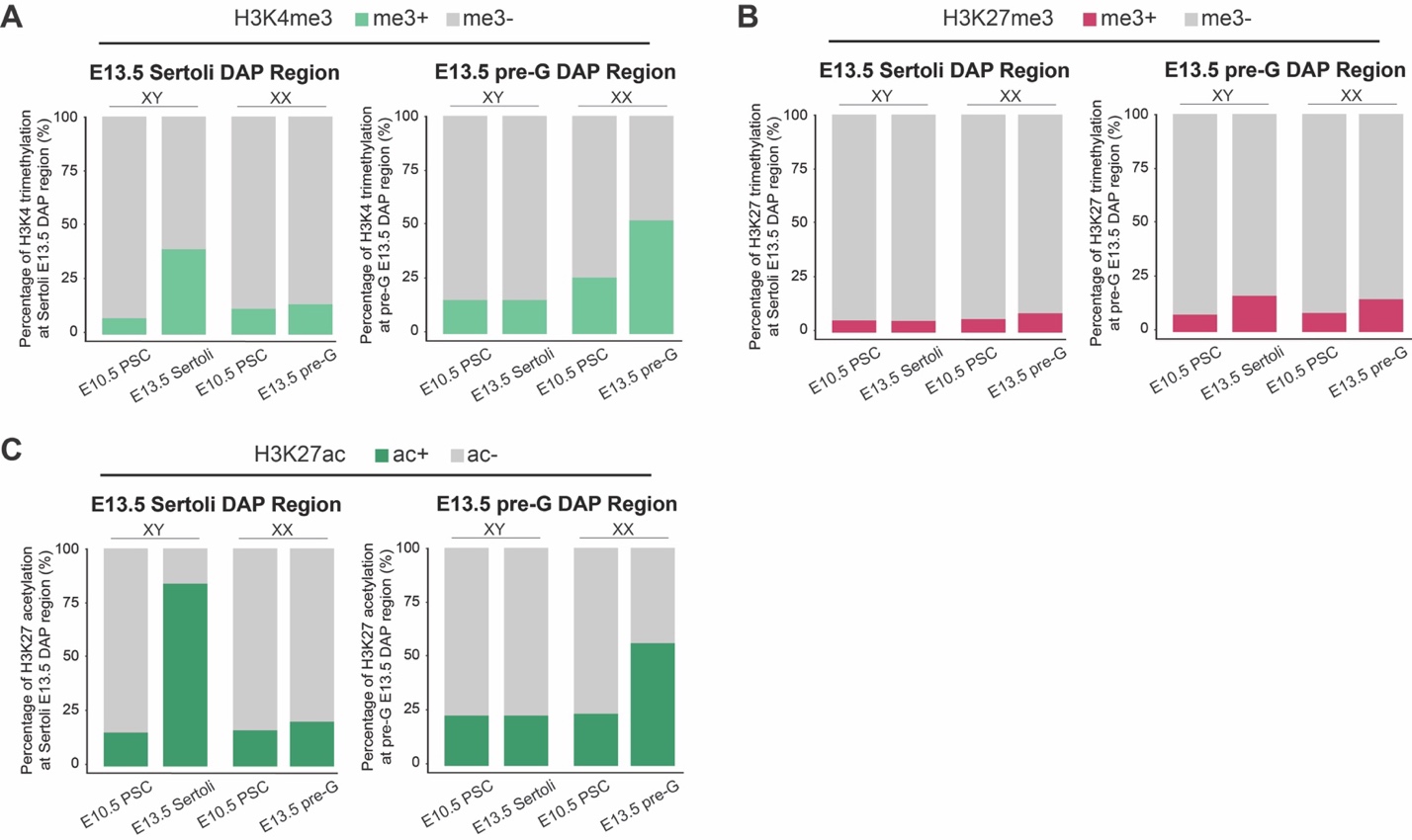

**Figure S4. Changes in histone modifications within E13.5 Sertoli and pre-granulosa DAPs.** (A-C) Percentage of chromatin regions positive for H3K4me3 (A), H3K27me3 (B), and H3K27ac (C), based on published ChIP-seq datasets (Garcia-Moreno et al, 2019), overlapping E13.5 Sertoli and pre-granulosa DAPs in XY and XX pre-supporting, Sertoli or pre-granulosa cells.

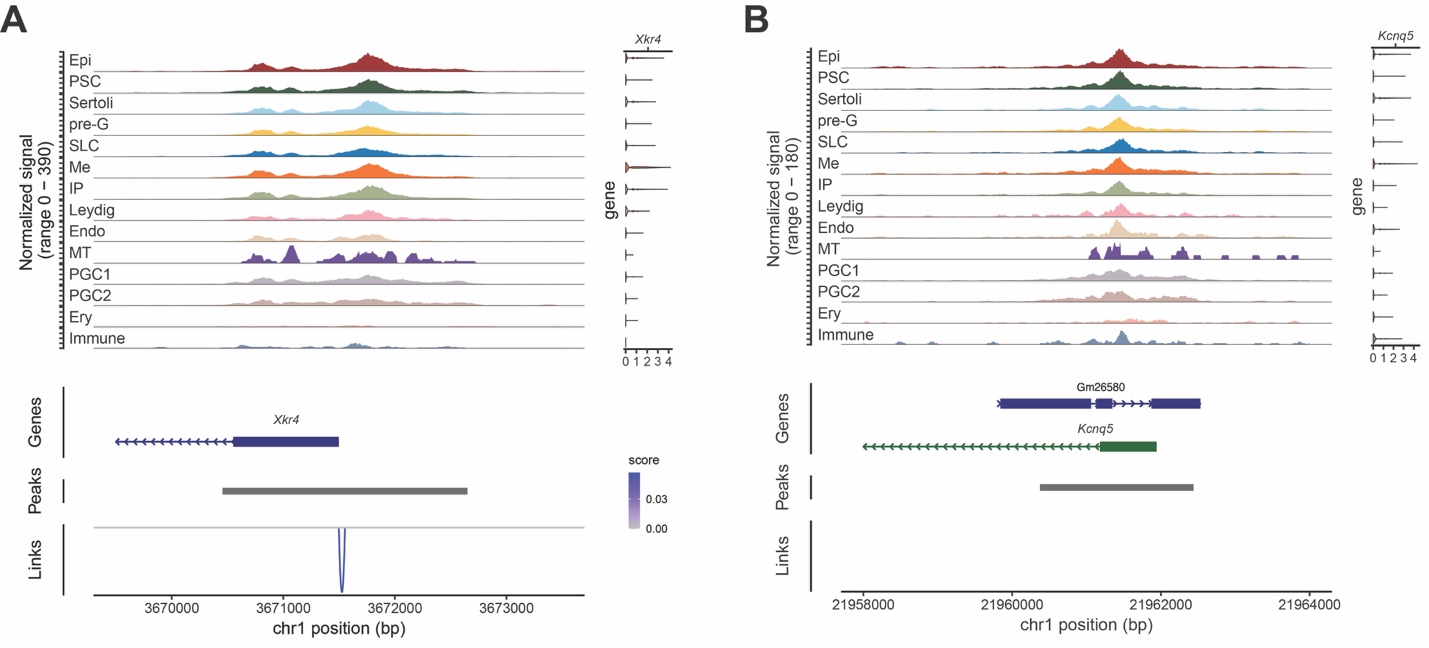

Figure S5. Linkage plot of Xkr4 and Kcnq5. (A) Peak-gene linkage plot for *Xkr4* across cell types from combined sex and developmental stages. The linkage score represents the level of association between chromatin accessibility and gene expression. (B) Peak-gene linkage plot for *Kcnq5* across cell types from combined sex and developmental stages, with no linkage identified. Annotations for the clusters are: Gonadal epithelial (Epi), Pre-supporting (PSC), Pre-granulosa (pre-G), Supporting-like cell (SLC), Mesenchymal (Me), Interstitial progenitor (IP), Endothelial cell (Endo), Mesonephric tubule (MT), Germ cell (PGC), Erythrocyte (Ery).

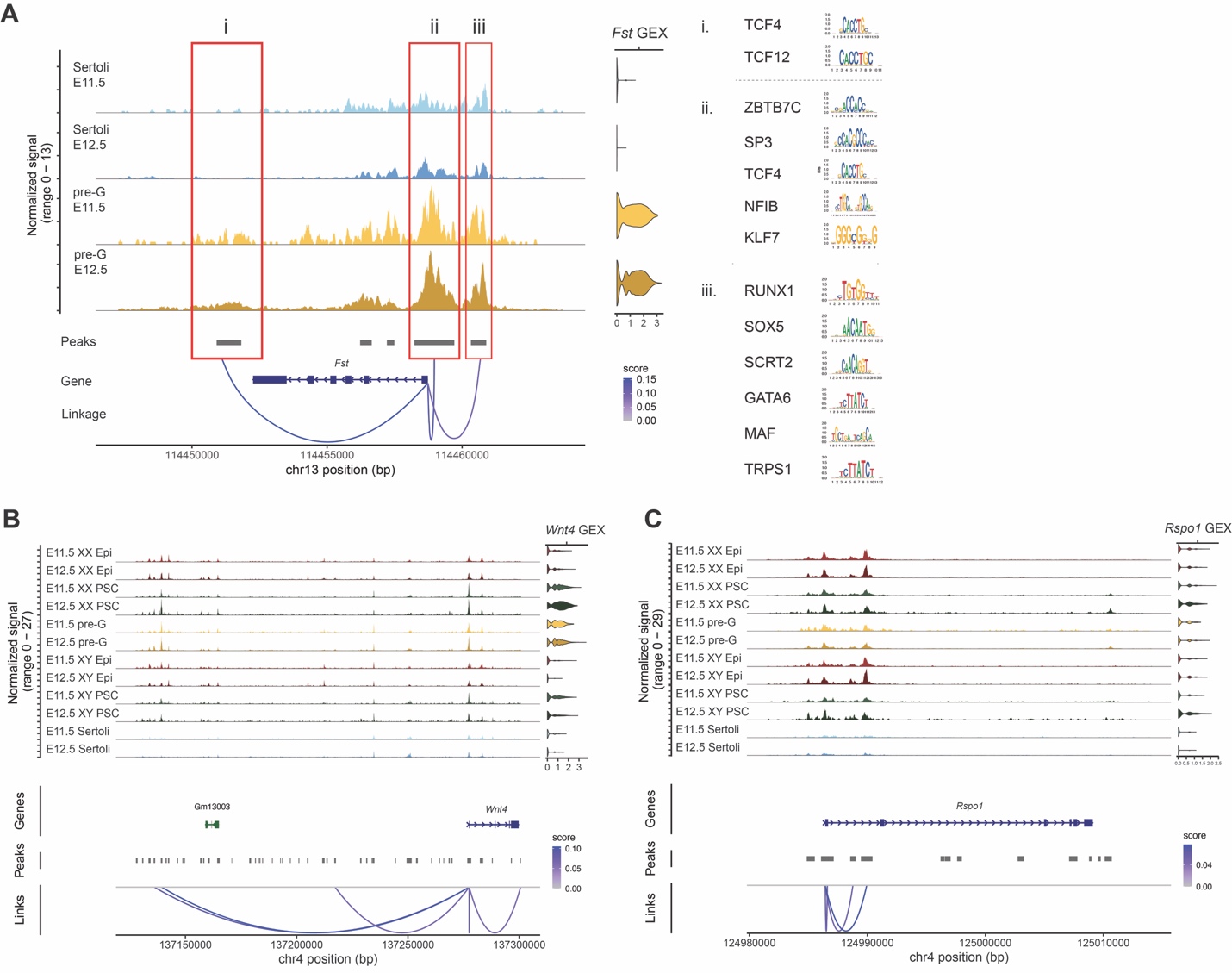

Figure S6. Linkage plots of *Fst*, *Wnt4*, and *Rspo*. (A) Pre-granulosa TF motifs identified within each peaks (i-iii) that are associated with *Fst* expression in E11.5 and E12.5 Sertoli and pre-granulosa cells. (B) Linkage plot of *Wnt4* in E11.5 and E12.5 XX and XY epithelial (Epi), pre-supporting (PSC), Sertoli, and pre-granulosa (pre-G) cells. (C) Linkage plot of *Rspo* in E11.5 and E12.5 XX and XY epithelial, pre-supporting, Sertoli, and pre-granulosa cells.

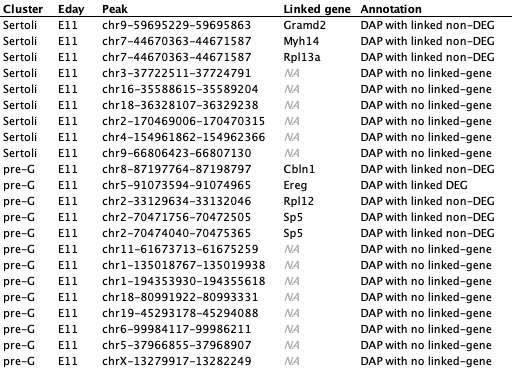

Table S1. Bivalent peaks within Sertoli and pre-granulosa E11.5 DAP regions. E11.5 Sertoli cell (Sertoli Cluster) or pre-granulosa cell (pre-G Cluster) specific DAPs (Peak) that are bivalent for both H3K4me3 and H3K27me3 histone marks. The table includes their associated genes (Linked gene) and whether these genes are differentially expressed between E11.5 Sertoli and E11.5 pre-granulosa cells (DEG versus non-DEG). Referenced in Figure 2D.

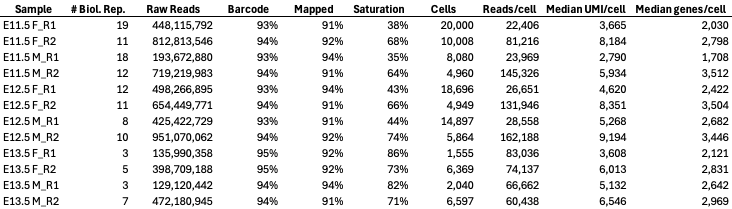

**Table S2.** **Sequencing and QC statistics**. Sequencing and QC statistics for single-nucleus multiome libraries of E11.5-E13.5 XX (F) and XY (M) gonads, including 3-19 biological replicates (pairs of gonads) and two technical replicates (R1 and R2).
